# Unbiased identification of *trans* regulators of ADAR and A-to-I RNA editing

**DOI:** 10.1101/631200

**Authors:** Emily C. Freund, Anne L. Sapiro, Qin Li, Sandra Linder, James J. Moresco, John R. Yates, Jin Billy Li

## Abstract

Adenosine-to-Inosine RNA editing is catalyzed by ADAR enzymes that deaminate adenosine to inosine. While many RNA editing sites are known, few *trans* regulators have been identified. We perform BioID followed by mass-spectrometry to identify *trans* regulators of ADAR1 and ADAR2 in HeLa and M17 neuroblastoma cells. We identify known and novel ADAR-interacting proteins. Using ENCODE data we validate and characterize a subset of the novel interactors as global or site-specific RNA editing regulators. Our set of novel *trans* regulators includes all four members of the DZF-domain-containing family of proteins: ILF3, ILF2, STRBP, and ZFR. We show that these proteins interact with each ADAR and modulate RNA editing levels. We find ILF3 is a global negative regulator of editing. This work demonstrates the broad roles RNA binding proteins play in regulating editing levels and establishes DZF-domain containing proteins as a group of highly influential RNA editing regulators.

## Introduction

RNA editing is a widely conserved and pervasive method of mRNA modification in which the sequence of a mRNA is altered from that encoded by the DNA (Nishikura, 2016; Walkley and Li, 2017). In mammals, the most prevalent type of RNA editing is Adenosine-to-Inosine (A-to-I) RNA editing (Eisenberg and Levanon, 2018). After editing occurs, inosine is recognized by the cellular machinery as guanosine (G); therefore, the editing of a nucleotide can have a variety of effects, including altering RNA processing, changing splice sites, and expanding the coding capacity of the genome (Burns et al., 1997; Nishikura, 2010; Rueter et al., 1999). A-to-I editing is catalyzed by adenosine deaminase acting on RNA (ADAR) proteins, which are conserved in metazoans (Nishikura, 2016).

Humans have two catalytically active ADAR proteins, ADAR1 and ADAR2, that together are responsible for millions of RNA editing events across the transcriptome. ADAR1 primarily edits long, near-perfect double-stranded RNA regions that are formed by inverted repeats, predominantly *Alu* elements (Athanasiadis et al., 2004; Bazak et al., 2014; Blow et al., 2004; Levanon et al., 2004). These editing events have been shown to play a role in self versus non-self RNA recognition in the innate immune response, and thus dysregulation of ADAR1 leads to immune-related diseases such as Aicardi-Goutieres syndrome (AGS) (Blango and Bass, 2016; Liddicoat et al., 2015; Mannion et al., 2014; Pestal et al., 2015; Rice et al., 2012). ADAR1 levels also correlate with tumor aggressiveness, as increases in ADAR1 editing suppress the innate immune response in tumors; accordingly, ADAR1 ablation helps with cancer therapy (Bhate et al., 2019; Gannon et al., 2018; Ishizuka et al., 2019; Liu et al., 2019; Nemlich et al., 2018). While the majority of ADAR1-regulated editing sites are found in repeat regions, ADAR2 is primarily responsible for editing adenosines found in non-repeat regions, particularly in the brain (Tan et al., 2017). ADAR2-regulated sites in non-repetitive regions include a number of editing events that alter the protein-coding sequences of neuronal RNAs, including *GluR2*, which encodes a glutamate receptor in which RNA editing is necessary for its proper function. Further demonstrating its important role in neuronal editing, dysregulation of human ADAR2 is associated with multiple neurological diseases, including amyotrophic lateral sclerosis, astrocytoma and transient forebrain ischemia (Slotkin and Nishikura, 2013; Tran et al., 2019). Maintaining RNA editing levels is critical for human health, but how RNA editing levels are regulated at specific editing sites across tissues and development is poorly understood.

While RNA sequence and structure are critical determinants of editing levels, studies querying tissue-specific and developmental-stage-specific editing levels show that the level of editing at the same editing site can vary greatly between individuals and tissues. These changes do not always correlate with ADAR mRNA or protein expression, suggesting a complex regulation of editing events by factors other than ADAR proteins (Sapiro et al., 2019; Tan et al., 2017; Wahlstedt et al., 2009). Recently, an analysis of proteins with an RNA-binding domain profiled by ENCODE suggested that RNA-binding proteins play a role in RNA editing regulation. Further, in mammals, a small number of *trans* regulators of editing have been identified through functional experiments (Quinones-Valdez et al., 2019). Some of these *trans* regulators of editing are site-specific, in that they affect editing levels at only a small subset of editing sites. These include RNA binding proteins such as DHX15, HNRNPA2/B1, RPS14, TDP-43, Drosha and Ro60 (Garncarz et al., 2013; Quinones-Valdez et al., 2019; Tariq et al., 2013). The recently identified ADAR binding partners, ELAVL1, DHX9 and SRSF9 have also been shown to affect the editing level of specific sites (Aktaş et al., 2017; Huang et al., 2018; Shanmugam et al., 2018; Stellos et al., 2016). In addition to site-specific regulators of editing, Pin1, WWP2 and AIMP2 have been shown to regulate editing through post-translational modification of the ADAR proteins (Behm et al., 2017; Marcucci et al., 2011; Tan et al., 2017). However, the complexity of editing level regulation across millions of editing sites in numerous tissues and developmental stages suggests that there are likely many other proteins that regulate editing.

Here, we take a unique approach to identify novel regulators of the ADAR proteins. We employ the BioID system, which facilitates the biotinylation and subsequent purification of proteins that both transiently and stably interact with bait proteins (Roux et al., 2012), to uncover proteins that interact with ADAR1 and ADAR2 in two human cell lines, HeLa and BE(2)-M17 cells. Together, these experiments facilitate the identification of 269 ADAR-interacting proteins, 15 of which had been previously reported, and many of which we further validate using publicly available data. Interestingly, the top candidates for novel regulators of ADARs represent a family of proteins that all contain a DZF-domain: ILF3, ILF2, STRBP, and ZFR. These proteins interact with both ADAR1 and ADAR2 in an RNA-dependent manner. We further characterize ILF3, the top candidate, and find that it acts as a negative regulator of editing that binds RNA near editing sites and globally regulates RNA editing levels. Furthermore, we demonstrate that the RNA binding domains of ILF3 are necessary for its editing regulation. This class of DZF-domain-containing proteins represents a novel group of RNA editing regulators and demonstrates the utility of the BioID experiment as a tool to systematically identify ADAR-interacting proteins that regulate RNA editing levels.

## Results

### ADAR1 and ADAR2 BioID identifies known and novel regulators of RNA editing

RNA editing levels are dynamically regulated, but few proteins responsible for this regulation are known. We set out to identify novel regulators of the ADAR proteins by identifying the proteins that ADAR1 and ADAR2 interact within the cell. We hypothesized that proteins that transiently interact with ADAR might compete with or recruit ADAR at only subsets of mRNA loci, making them good candidates for site-specific regulators of RNA editing. To identify these proteins, we utilized the BioID system (Roux et al., 2012). This method expands upon traditional immunoprecipitation (IP) and mass-spectrometry-based screening used to identify protein-protein interactions. Traditional IPs enrich for proteins that form a stable complex with the bait protein but are less effective at identifying transient interactions. The BioID protocol efficiently captures both this transiently interacting class of regulators and the more standard stable protein partners by covalently labeling factors as they come into close proximity (~10 nm) with a bait protein (Kim et al., 2014). These labeled proteins can then be immunoprecipitated and identified by mass-spectrometry (**Figure 1A**). To utilize this system, we fused ADAR proteins with a mutated form of the BirA biotin ligase (R118G), denoted BirA* (Kwon et al., 2000; Roux et al., 2012), which promiscuously and irreversibly biotinylates proteins in a proximity-dependent manner.

**Figure 1.**
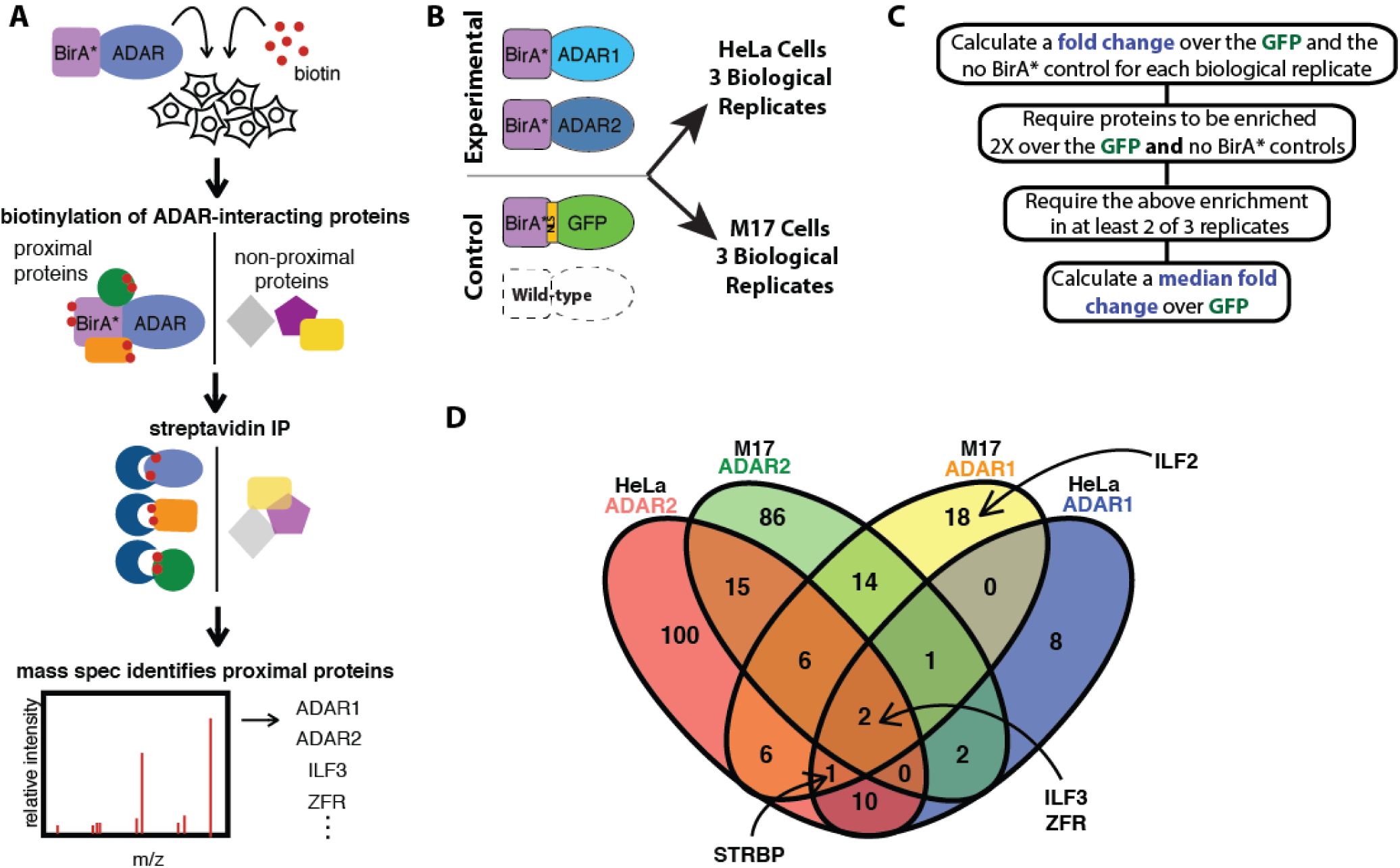
BioID in human cells identifies known and novel regulators of ADAR1 and ADAR2. **A.** Schematic depicting BioID protocol for ADAR. A fusion of ADAR1 or ADAR2 with the BirA* enzyme was expressed in cells and the media was supplemented with exogenous Biotin, allowing for the fusion protein to biotinylate proximal proteins. A Streptavidin IP was then performed to isolate biotinylated proteins. The eluted proteins were then identified by mass spectrometry. **B.** Schematic of BioID experimental conditions. The experiment was performed with two different BioID fusion proteins, BirA*-ADAR1 and BirA*-ADAR2, and two negative controls, BirA*-GFP and untransfected cells with no BirA expressed. Each fusion protein and control was assayed in HeLa and M17 cells for a total of eight experimental conditions, performed in triplicate. **C.** Pipeline depicting the analysis performed to determine and rank BioID hits. A series of filters were applied using each control condition to remove false positives. Median fold change over the GFP condition was used to rank the hits. **D**. Venn diagram depicting the hits from each BioID experiment. The hits from each BioID are indicated by differently colored ovals (pink: HeLa ADAR2, green: M17 ADAR2, Yellow: M17 ADAR1, Blue: HeLa ADAR1). The numbers indicate the number of hits that overlap between each condition and the proteins denoted in bold are the BioID hits further characterized in this study.

We lentivirally integrated either BirA*-ADAR1 or BirA*-ADAR2 constructs into two human cell lines, neuroblastoma BE(2)-M17 (M17) cells and HeLa cells. By using two cell lines, we hoped to identify both tissue-specific and universal editing regulators. To first demonstrate that the constructs were functional, we assessed editing levels at two editing sites. We observed an increase in editing at an ADAR1-specific site in *PAICS* upon expression of BirA*-ADAR1 and an increase in editing at the ADAR2-specific site in *GRIA2* upon expression of BirA*-ADAR2 compared to a BirA*-GFP control (**Figure S1A**), demonstrating that the BirA*-ADAR constructs produce functional ADAR proteins.

After verifying the activity of the BirA*-ADAR proteins, we performed the BioID experiments. We produced three biological replicates for both BirA*-ADAR1 and BirA*-ADAR2 in both M17 and HeLa cells, along with three replicates of two different negative controls: nuclear localized BirA*-GFP and cells with no transgene and thus no BirA* expression in each cell line (**Figure 1B**). The cells were incubated overnight with D-biotin to allow the fusion proteins to biotinylate interactors. We then performed a stringent denaturing pulldown with streptavidin agarose and analyzed the elution by mass spectrometry (see **Methods** for details). To identify the putative ADAR1- and ADAR2-interacting proteins from our list of all proteins returned from the mass-spectrometry, we determined the fold change in peptide counts for each protein in each BirA*-ADAR condition compared to BirA*-GFP and the no BirA* controls, then defined hits as those proteins having at least two-fold enrichment over both of the negative control conditions in at least two biological replicates (**Figure 1C, S3A**). In total, we identified 269 proteins as putative interactors of either ADAR1 and/or ADAR2 in HeLa and/or M17 cells. When comparing across all 4 conditions, we found hits unique to each condition: 26 proteins unique to ADAR1 and 201 unique to ADAR2, 127 proteins unique to Helas and 118 unique to M17 cells, and 57 proteins shared across multiple conditions (**Figure 1D, S3B,C**). Supporting the efficacy of this screening approach, we found many of the proteins previously reported to affect editing or interact with ADAR proteins. The ADAR-interacting proteins we identified included DHX15, RPS14, and ELAVL1, which were previously reported to regulate editing at specific sites, and SFPQ, HNRNPH1 and PTBP1, which are known post-transcriptional regulators of ADARs (Hirose et al., 2014; Yang et al., 2015). We also identified known ADAR-interacting protein CPSF6 and proteins involved in L1 Line element retrotransposition that were previously found in a complex with ADAR1: DHX15, SFPQ, NCL, TUBB, NONO, HSPA8, SF3B1, and HNRNPL (Binothman et al., 2017; Orecchini et al., 2017). Furthermore, we identified proteins that, like ADAR, bind the *ACA11* transcript: SF3B1, PARP1, NCL, DDX21 and ILF3 (**Figure 1D, S3C**) (Chu et al., 2012).

Overall, we found many more candidates specific to ADAR2 than ADAR1, which is likely due to the fact that ADAR2-BirA* was more highly expressed than ADAR1-BirA*; we found that overexpressed ADAR2 consistently accumulates to higher protein levels than overexpressed ADAR1, perhaps reflecting different mechanisms of regulation (data not shown). We also found a large number of cell-type-specific ADAR interactors (118 M17 versus 127 HeLa hits), which we hypothesized might be differentially expressed between the two cell types. To determine whether cell-type-specific interactors were more highly expressed in the cell type in which they were identified, we measured the expression level of our candidates in each cell line using RNA-seq. We found that candidates specific to M17 cells were more highly expressed in M17 cells than in HeLa cells (as measured by FPKM, Wilcoxon test, p < 0.001), but the reverse was not true, suggesting that differential mRNA expression does not fully explain the specificity of proteins interacting with ADARs in these two cell types (**Figure S3A,B**).

### Novel ADAR-interacting proteins bind near editing sites to alter editing levels

Many proteins found in the BioID screen were RNA binding proteins and RNA processing enzymes, suggesting enrichment for biologically relevant proteins. We utilized publicly available RNA-seq data from shRNA knockdowns performed in K562 cells by the ENCODE project (Sundararaman et al., 2016) to identify whether any of the RNA binding proteins identified in the BioID altered editing levels. We found that knockdown of 8 of the 19 proteins profiled by ENCODE resulted in increases or decreases in the editing levels of more than 100 sites, while the remaining 11 affected a smaller more specific set of sites (**Figure 2A, 2S**). We hypothesized that RNA-binding proteins that affected editing might bind RNA near editing sites and alter ADAR binding and therefore editing levels at those sites. To test whether this was true of our 19 candidate interactors, we utilized the publicly available eCLIP-seq data (“A Large-Scale Binding and Functional Map of Human RNA Binding Proteins | bioRxiv,” n.d.) to identify the RNA binding sites of the 19 candidates profiled by ENCODE. We determined the proximity of each protein’s RNA binding sites to known editing sites. Of these 19 candidates, 4 showed evidence of binding at editing sites (**Figure 2B**) and 9 showed evidence of binding nearby, but not right at, editing sites (**Figure 2C**). This is consistent with the hypothesis that these RBPs might interfere with or recruit ADARs to these editing sites. The remaining 6 candidates did not show evidence of binding near editing sites (**Figure 2D**). Included in this last group is PTBP1, which has been shown to be a translational regulator of ADAR (Yang et al., 2015); this role may explain why it physically interacts with ADAR but does not bind near editing sites. For the 13 RBPs that bound RNA at or near editing sites, we compared the positions of their binding sites to the editing sites that they regulated to determine whether their knockdown affected editing levels specifically at the sites that were bound by the proteins. We found a statistical enrichment for 11 of the 13 candidates indicating that they changed editing levels at editing sites near where they bound RNA, suggesting a direct role in RNA editing (**Figure 2E**). Of those proteins, only U2AF2 and XRCC6 altered the majority of known editing sites in a single direction (**Figure 2A,E**), suggesting that RBPs binding at or near editing sites does not necessarily lead to the same changes in editing levels at every site.

**Figure 2.**
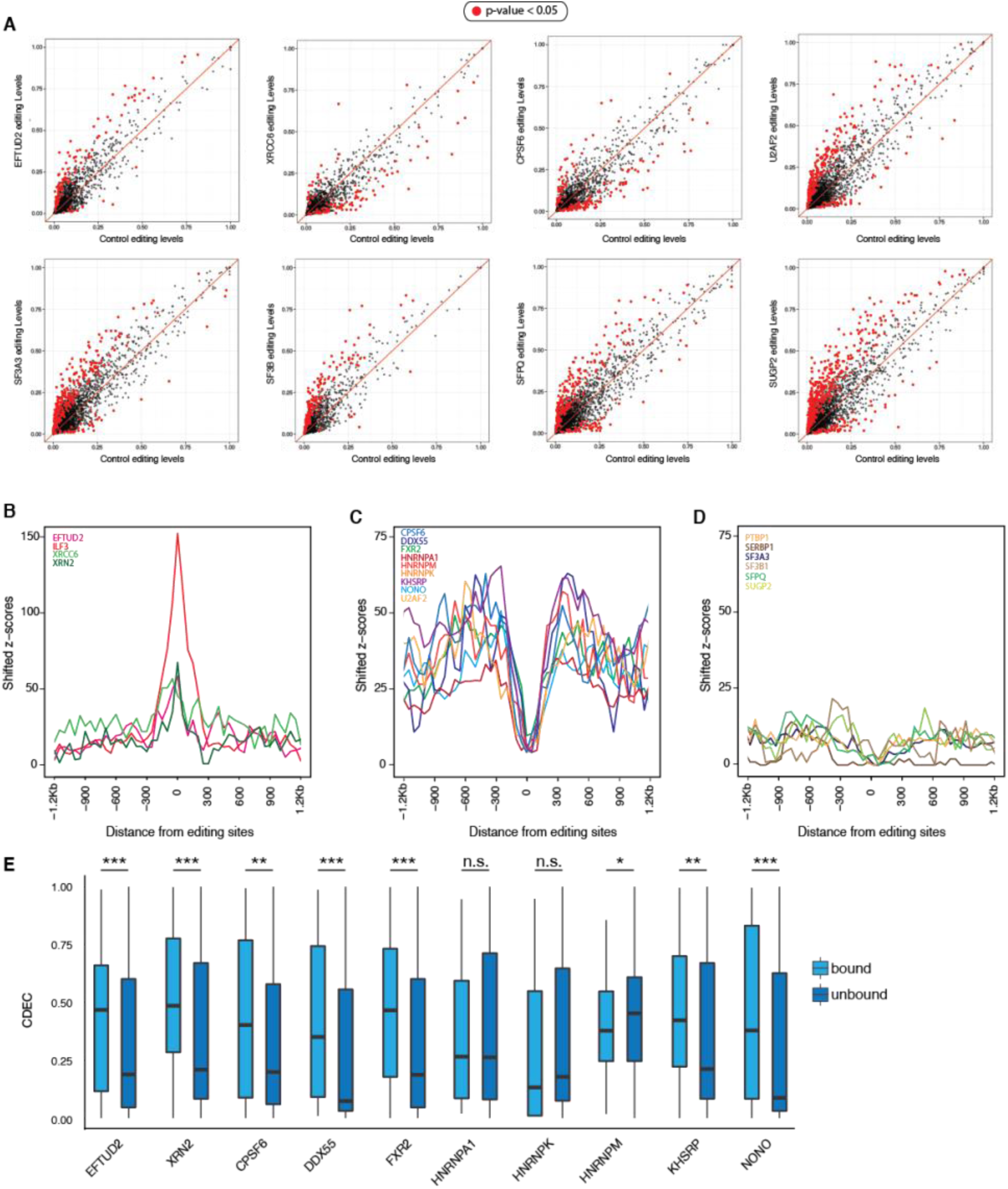
RNA-seq and eCLIP-seq analyses identify regulatory RNA-binding proteins of RNA editing. **A.** Scatter plots of pairwise comparison of editing levels between knockdown and control RNA-seq of 8 top regulatory RBPs. Red dots, Fisher’s exact test p-value < 0.05. **B-D**. Z-scores showing the observed RBP binding strength over the expected one, measured in standard deviations (see **Methods** section for details). **B.** RBPs that bind directly to editing sites in eCLIP-seq. **C.** RBPs that bind near-by editing sites (non-overlapping) in eCLIP-seq. **D.** RBPs that show no enrichment of binding near editing sites in eCLIP-seq. **E.** Comparison of editing profile differences in knockdowns between sites bound by RBPs (light blue) and the ones unbound (dark blue). CDED: <u>c</u>umulative <u>d</u>istribution of <u>e</u>diting level <u>d</u>eviation, quantifies the accumulative editing level difference from the mean between controls and knockdowns (**Methods**).

### DZF-domain-containing proteins interact with ADAR1 and ADAR2 in an RNA-dependent manner

One group of proteins that we identified in the BioID were four related proteins, ILF2, ILF3, STRBP and ZFR, which are the only four human proteins that contain a DZF domain. The DZF domain (domain associated with zinc fingers) is a poorly understood domain, but it has been shown to drive protein dimerization and facilitate RNA binding (Castello et al., 2016; Wolkowicz and Cook, 2012). Two of these DZF-domain-containing proteins, ILF3 and ZFR, were the only proteins identified as hits in all four BioID conditions (ADAR1- and ADAR2-interactors in both cell types). ILF3 has been shown previously to interact with ADAR1-p150 in an RNA-dependent manner in the cytoplasm (Nie et al., 2005). Our results extend its interaction to ADAR-p110, the isoform we overexpressed in our BioID experiments. Another DZF-domain-containing protein, STRBP, was identified in three out of four conditions (all except HeLa BirA*ADAR1) and the fourth, ILF2, was identified in the M17 BirA*-ADAR1 condition, strongly implicating DZF-domain-containing proteins as ADAR-interacting proteins. In addition to a DZF domain, all three proteins except for ILF2 contain double-stranded RNA binding domains (dsRBDs), similar to ADAR1 and ADAR2 (**Figure 3A**). ILF3 and STRBP have two dsRBDs, while ZFR has three widely spaced zinc finger domains, which are thought to mediate interaction with dsRNAs. Intriguingly, ILF3’s dsRBDs have been shown to be structurally similar to those of ADAR2 and to compete for similar binding sites *in vitro* (Wolkowicz et al 2012). In addition, ILF3 has been shown to regulate circular RNA biogenesis by binding near the highly edited *Alu* elements (Li et al., 2017). Because this entire class of proteins was highly enriched in the BioID hit list and had been previously suggested to interact with ADAR1, we chose to further characterize this class of proteins to understand their regulation of RNA editing.

**Figure 3.**
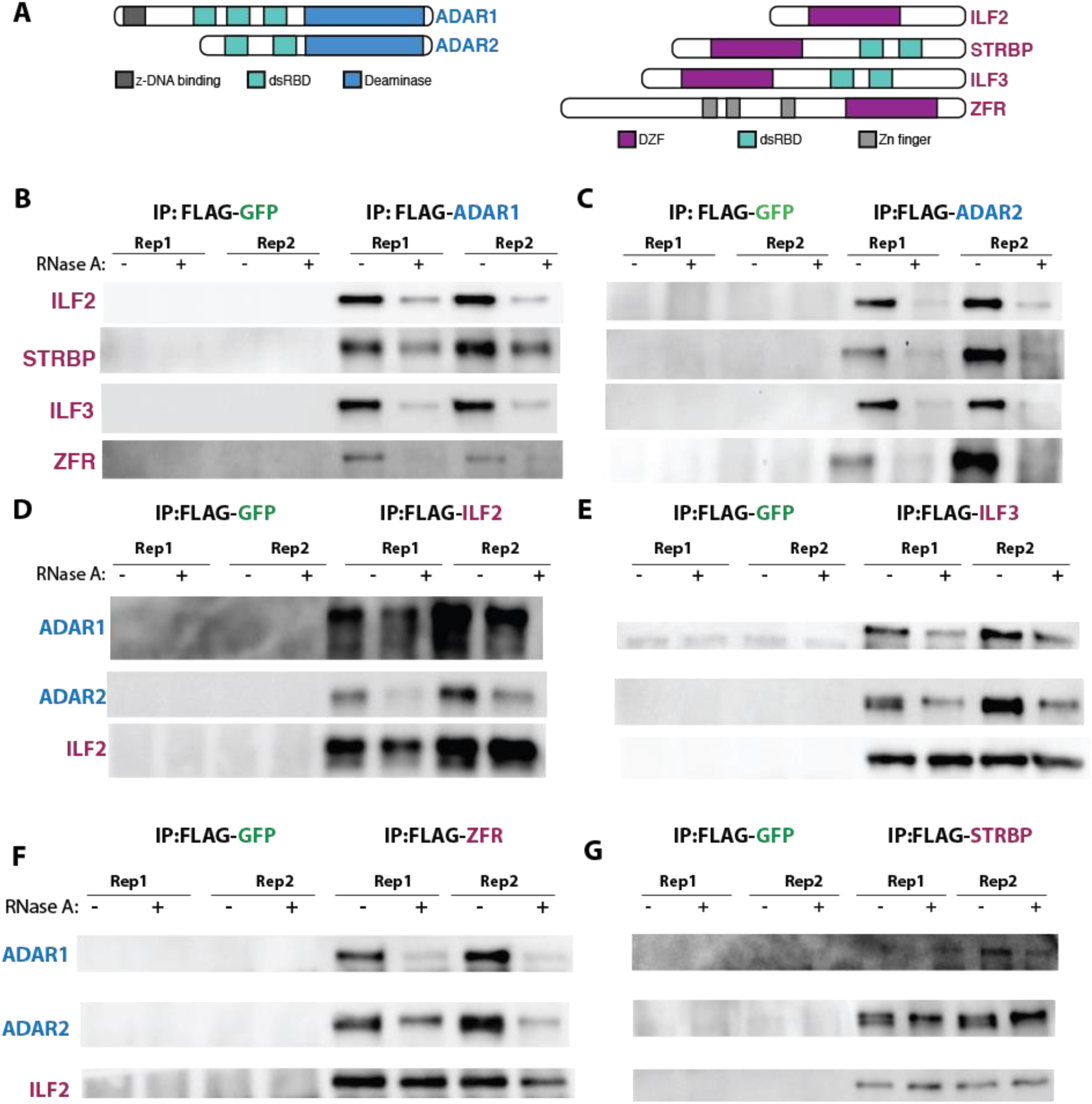
co-IPs demonstrate that DZF-domain-containing BioID hits interact with ADAR1 and ADAR2 in an RNA-dependent manner. **A.** Schematic depicting ADAR1, ADAR2 and DZF-domain-containing proteins ILF2, STRBP, ILF3, and ZFR. Conserved domains are indicated with different colors. **B-C.** Western blots of FLAG immunoprecipitation of M17 cells overexpressing either FLAG-GFP (negative control), FLAG-ADAR1, or FLAG-ADAR2. IPs were performed with or without the addition of RNase A to the lysates prior to IP, as indicated. **D-G.** Western blots of FLAG immunoprecipitation of M17 cells overexpressing ADAR2 as well as FLAG-GFP (negative control) or a FLAG tagged version of each DZF-domain-containing protein, (D) ILF2, (E) ILF3, (F) ZFR, (G) STRBP. IPs were performed with or without the addition of RNase A to the lysates prior to IP, as indicated. ILF2, which interacts with itself, ILF3, ZFR, and STRBP in an RNA-independent manner, was used as a positive control.

To validate the interaction between ILF3, ZFR, STRBP, and ILF2 and each ADAR protein, we performed traditional co-IPs in M17 cells. We transduced a FLAG-tagged ADAR1, ADAR2, or GFP as a negative control, into M17 cells (**Figure S5A, B**) and performed an IP using an antibody against FLAG. To further determine whether the interaction between each ADAR and each DZF-domain-containing candidate was dependent on RNA, we treated half of each of the IPs with RNase A. Whereas FLAG-GFP did not immunoprecipitate any of the four candidates, FLAG-ADAR1 and FLAG-ADAR2 immunoprecipitated each candidate. In all cases, the interaction between each candidate and each ADAR was decreased upon addition of RNase A, suggesting that their interaction is at least partially RNA-dependent, and that they do not form a stable complex off of RNA (**Figure 3B, C**).

We also wanted to interrogate the reciprocal condition, the ability of each candidate to immunoprecipitate each ADAR protein; however, M17 and HeLa cells do not express high levels of ADAR2, making it difficult to analyze endogenous ADAR2’s interaction with each candidate in these cells. To address this problem, we created an M17 cell line overexpressing ADAR2 by lentiviral transduction. We then stably expressed a FLAG-tagged version of each candidate, or FLAG-GFP as a negative control, in the ADAR2-overexpressing M17 cell line (M17-ADAR2-OE) (**Figure S5C-F**). We performed FLAG IPs with and without RNase A treatment in this cell line. We used pulldown of ILF2 as a positive control because previous work established that it binds each of the other DZF-domain-containing proteins in an RNA-independent manner, and thus we would expect to see that it binds each candidate protein with and without RNase A (Wolkowicz and Cook, 2012). As expected, ILF2, ILF3, and ZFR all bound ADAR1 and ADAR2 in an RNA-dependent manner and bound ILF2 independent of RNA (**Figure 3D-G**). Unexpectedly, in contrast to the FLAG-ADAR2 IP, in the FLAG-STRBP IP we found an RNA-independent interaction between STRBP with ADAR2 (**Figure 3G**), which could indicate a different mode of interaction depending on the bait protein. Taken together, we were able to fully validate that three of our top candidates biochemically interact with ADARs in an RNA-dependent manner.

### DZF-domain-containing proteins affect A-to-I RNA editing

Similar to the validation of other candidates that we performed with publicly available RNA-seq and eCLIP-seq data, we wanted to perform a more thorough analysis of the role of DZF-domain-containing proteins in A-to-I RNA editing. We transiently overexpressed ILF2, ILF3, and STRBP in HEK293T cells and in HEK293T cells stably overexpressing ADAR2 through lentiviral transduction (HEK293T-ADAR2-OE) (**Figure 4A, B**); as in M17 cells, ADAR2 is lowly expressed in HEK293T cells, thus requiring overexpression for analysis of ADAR2-controlled editing sites. We choose to perform these experiments in HEK293T cells both because in contrast to M17 cells, they were suitable for transient transfection and it extended our findings to a third cell type. We first verified that each protein was overexpressed at the transcript level (**Figure 4A, B**) and protein level **(Figure S6A,B**). To examine the effects of the DZF-domain-containing proteins on editing levels, we used microfluidic multiplex PCR and sequencing (mmPCR-seq), in which we PCR-amplified cDNA at thousands of known editing sites for subsequent Illumina sequencing [REF]. mmPCR-seq editing level measurements were highly reproducible between two biological replicates from each cell type (**Figure S7A, Table S1**). We compared the editing levels at these highly reproducible sites between the GFP-overexpressing control and the cells overexpressing DZF-domain-containing proteins in both HEK293T and HEK293T-ADAR2-OE backgrounds. Overexpression of ILF2 had a moderate effect in both HEK293T and HEK293T-ADAR2-OE cells (**Figure 4C, D**); 11 sites had reduced editing in HEK293T cells, while 15 sites showed increased editing, and 18 sites showed reduced editing in HEK293T-ADAR2-OE cells. This relatively weak and bidirectional effect on editing is consistent with ILF2’s identification in only one of the four BioID conditions (interacting with ADAR1 in M17 cells), suggesting that on its own it may not be a robust ADAR interactor. STRBP demonstrated a slightly stronger effect; 19 sites were reduced in HEK293T cells and 27 in HEK293T-ADAR2-OE cells with only 7 and 3 sites increased in editing respectively (**Figure 4E, F**). By far, ILF3 demonstrated the strongest effect; 39 sites were decreased in HEK293T cells and 47 in HEK293T-ADAR2-OE cells with only 4 and 1 editing sites showing increased editing levels, respectively (**Figure 4G, H**). To better understand the role of ILF3 in editing regulation, we performed RNA-seq on cells with ILF3 overexpression in both HEK293T and HEK293T-ADAR2-OE backgrounds. Similar to mmPCR-seq, editing level measurements were highly reproducible between two biological replicates from each cell type (**Figures S7B, Table S2**). This more expansive genome-wide analysis further revealed ILF3 to be a global regulator of editing, as almost the entire editing landscape was reduced when ILF3 was overexpressed. We observed 221 sites with a significantly reduced editing level (p < 0.05, Fisher’s exact test) compared to only 16 with an increased editing level in HEK293T cells, and we found 237 sites with a significantly reduced editing level compared to only 2 with an increased editing level in HEK293T-ADAR2-OE cells (**Figure 4I**). In addition, we calculated overall editing as the total number of edited G reads over the total of A + G reads at all known editing sites combined as a global measure of ADAR activity (Tan et al 2017). We found that in both HEK293T and HEK293T-ADAR2-OE cells, overall editing was significantly reduced, from 2.43% to 1.45% in HEK293T cells, and from 4.56% to 2.45% in HEK293T-ADAR2-OE cells (**Figure 4J**). Overall, our data show that ILF3 and STRBP are both negative regulators of editing but that ILF3 is a stronger global regulator of editing.

**Figure 4.**
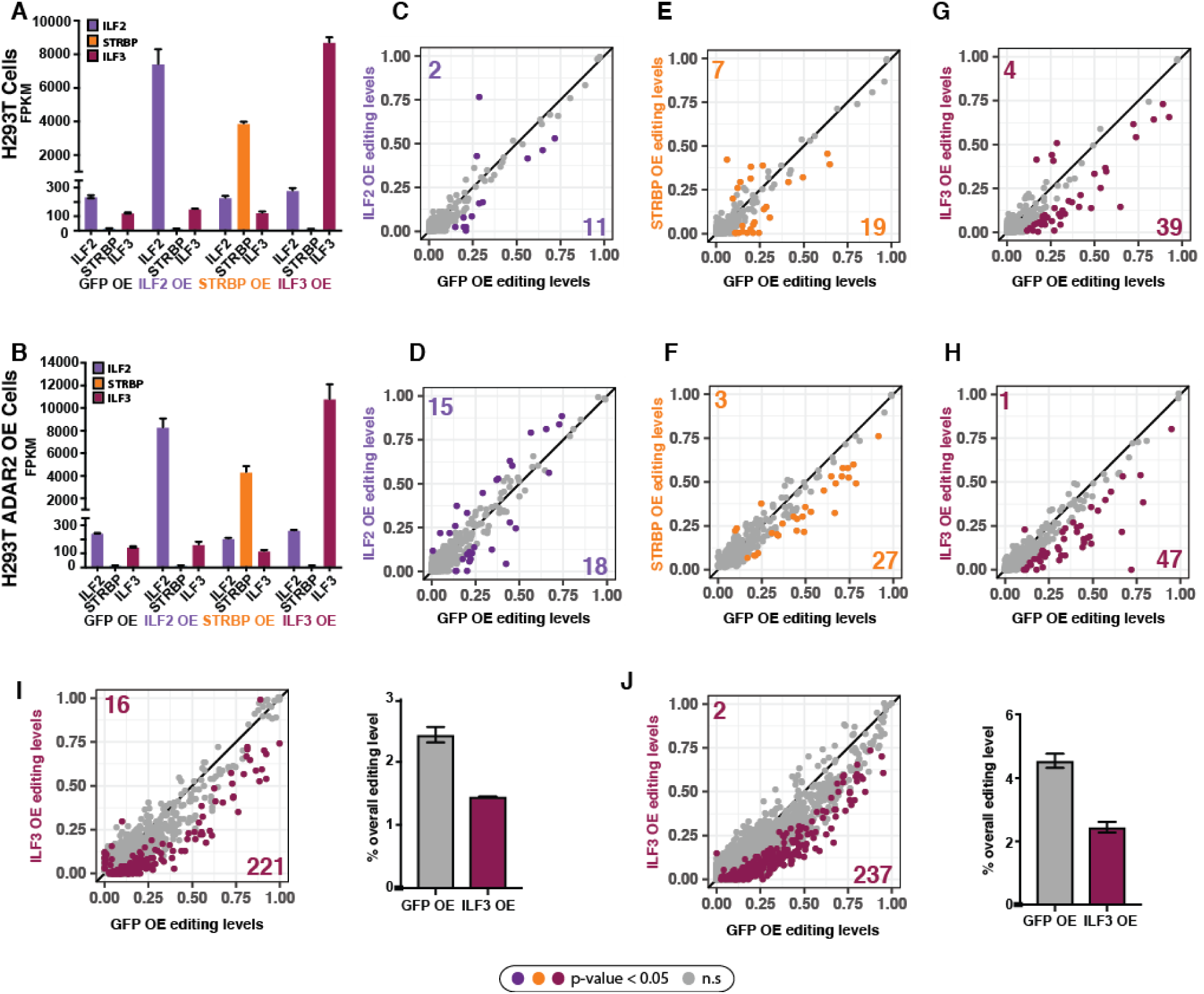
Overexpression of DZF-domain-containing proteins alters RNA editing levels. **A-B.** FPKMs of ILF2, STRBP, and ILF3 in (A) HEK293T and (B) HEK293T-ADAR2-overpression cells transiently transfected with GFP, ILF2, STRBP, or ILF3. Error bars represent standard deviation. **C-H.** Scatterplots comparing RNA editing levels assayed by mmPCR-seq in HEK293T cells (upper) and HEK293T cells stably overexpressing ADAR2 (lower). Plots compare editing levels in cells overexpressing GFP with editing levels in cells overexpressing a DZF-domain-containing protein ILF2 (C,D), STRBP (E,F) ILF3 (G,H). Colored dots indicate sites that are significantly changed (p < 0.05, Fisher’s exact tests). The number of sites with significantly increased (top left) or significantly decreased (bottom right) editing levels are indicated on each graph. **I-J.** Scatterplots comparing RNA editing levels assayed by RNA-seq in HEK293T cells (I) and HEK293T cells stably overexpressing ADAR2 (J). Plot compares editing levels in cells overexpressing GFP with editing levels in cells overexpressing ILF3. Colored dots indicate sites that are significantly changed (p < 0.05, Fisher’s exact tests). The number of sites with significantly increased (top left) or significantly decreased (bottom right) editing levels are indicated on each graph. Bar graphs to the right of each scatterplot depict overall editing levels (percentage of edited reads at all sites) for GFP overexpression (grey) and ILF3 overexpression (red).

One possible explanation for the effect of DZF-domain-containing proteins on RNA editing could be transcriptional or translational regulation of ADAR mRNA or protein levels. To test this hypothesis, we analyzed the mRNA and protein levels of ADAR1 and ADAR2 in ILF2-, ILF3-, and STRBP-overexpressing cells compared to GFP-overexpressing cells in HEK293T and HEK293T-ADAR2-OE cells. We found that ADAR1 mRNA and protein levels were largely unchanged (**Figure S7A, B**). *ADAR2* transcript levels were reduced, which may account for some of the editing effects in HEK293T-ADAR2-OE cells (**Figure S8A, B**); however, there was a reduction in editing even in HEK293T cells, which largely lack ADAR2 expression and ILF2 overexpression showed a similar reduction in ADAR2 protein without a strong effect on editing, suggesting regulation of ADAR2 levels cannot be the sole mechanism that DZF-domain-containing proteins use to repress editing.

### RNA binding activity of ILF3 is necessary for its regulation of RNA editing

Because changes in ADAR levels did not explain the differences in editing that we observed, we hypothesized that the global regulator ILF3 and ADAR proteins compete for the same transcripts. This hypothesis is supported by previous work showing that ILF3 has structurally similar dsRBDs to ADAR2 (Wolkowicz and Cook, 2012). This mechanism would be consistent with our previously observed RNA-dependent interaction between ILF3 and the ADARs. In addition, we and others have shown that ILF3 binds near editing sites in K562 cells (**Figure 2B**) (Quinones-Valdez et al., 2019). If ILF3 competes for RNA binding sites with ADAR proteins, then the ability of ILF3 to affect editing levels would be dependent on its ability to bind RNA. To test this hypothesis, we overexpressed a FLAG-tagged mutant of ILF3 that lacks its two dsRBDs (Δ402-572) in HEK293T and HEK293T-ADAR2-OE cells (**Figure 5A, B, S9A, Table S1**). In both cases, ILF3-ΔdsRBD-OE did not induce a global reduction in editing (**Figure 5C, S9B**), suggesting that ILF3’s RNA binding ability is necessary for its regulation of editing levels. In addition, we performed a FLAG IP to pull down the mutant and wild-type version of ILF3 and found that only wild-type ILF3 interacts with ADAR1 and ADAR2 (**Figure 5D**). This result further supports the hypothesis that it is the ability of ILF3 to bind RNA and compete for substrates with ADARs that regulates editing levels.

**Figure 5.**
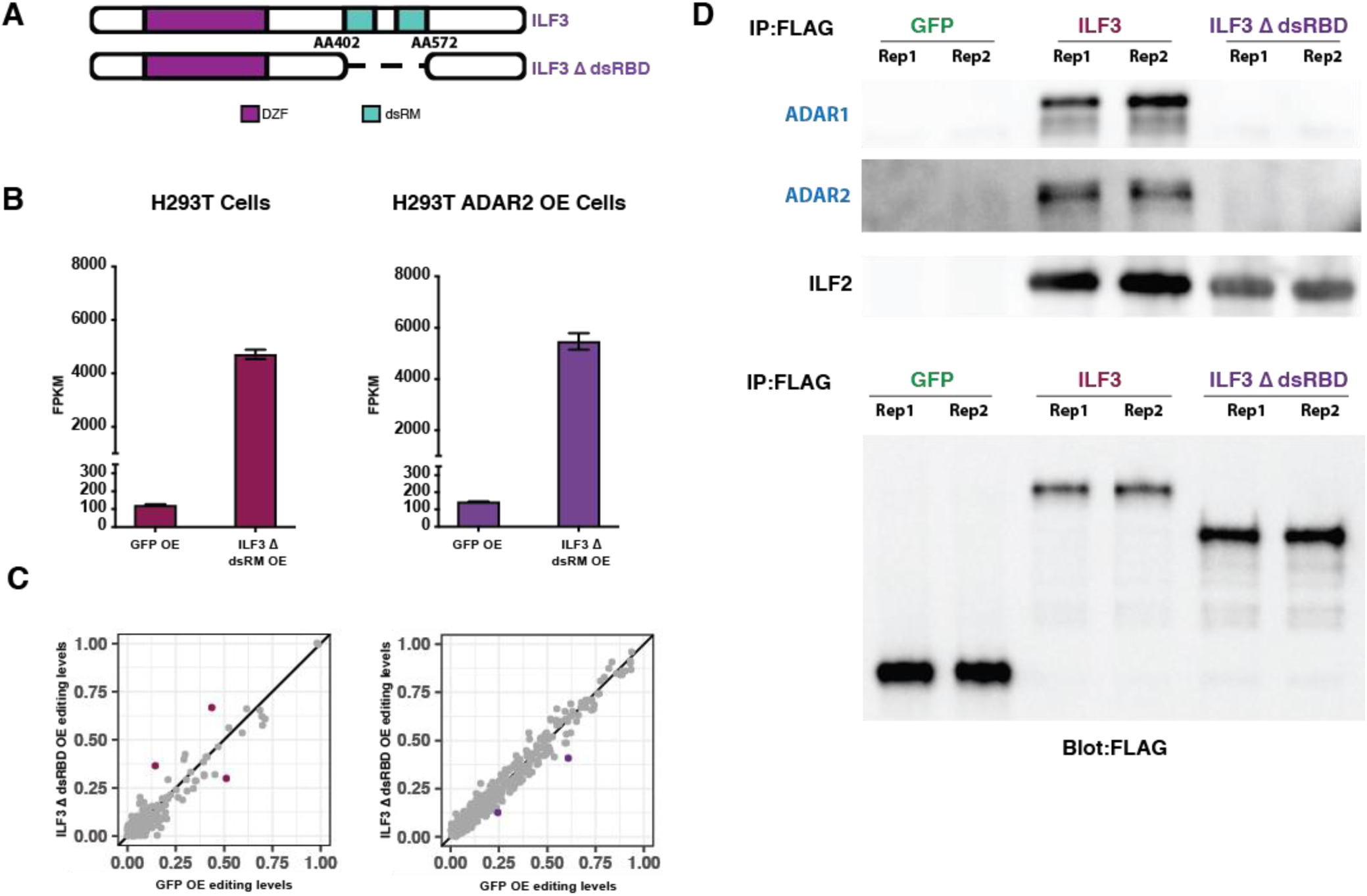
Overexpression of ILF3 dsRNA-binding mutant does not interact with either ADAR or affect editing levels. **A.** Schematic of ILF3 double-stranded RNA binding domain (dsRBD mutant, dashed lines indicate the deleted region (which spans amino acids 402-572). **B.** Bar graphs depicting the FPKM of ILF3 in GFP and ILF3ΔdsRBD overexpression in HEK293T cells (left) and HEK293T cells stably overexpressing ADAR2 (right). **C.** Scatterplots comparing RNA editing levels assayed by mmPCR-seq in HEK293T cells (left) and HEK293T cells stably overexpressing ADAR2 (right). Plot compares editing levels in cells overexpressing GFP (negative control, x-axis) with editing levels in cells overexpressing ILF3ΔdsRBD (y-axis). Colored dots indicate sites that are significantly changed (p < 0.05, Fisher’s exact tests). Overexpression of the ILF3 dsRBD mutant does not affect editing levels. **D.** Western blots of FLAG immunoprecipitation of HEK293T cells overexpressing ADAR2 and either FLAG-GFP (negative control), FLAG-ILF3, or FLAG-ILF3ΔdsRBD. ILF3ΔdsRBD mutant does not interact with ADAR1 or ADAR2, but maintains its interaction with ILF2.

## Discussion

RNA editing is widely conserved and pervasive, leading to changes at the RNA level. It is catalyzed by two enzymatically active ADARs, ADAR1 and ADAR2. Expression levels of these ADAR enzymes do not correlate with the editing frequency of large classes of sites (Tan et al., 2017), which strongly suggests the presence of additional editing regulators. We performed a large-scale, unbiased assay to systematically identify novel editing regulators in humans by employing a biochemical-based screening approach, BioID. BioID labels proteins in proximity of a BirA*-tagged bait protein, thus allowing us to identifying proteins that physically interact with ADARs transiently or stably. As editing regulation is known to be cell-type-specific, we performed the BioID screen in two different cell types, HeLa and M17, using either ADAR1 or ADAR2 as bait. This approach was highly successful in enriching for ADAR interactors, in that it identified most previously known ADAR binding partners and editing regulators. The small number of previously known ADAR-interactors that were not found here probably interact with ADAR proteins in a cell-type-specific context that may not exist in our system. While many proteins were found in multiple BioID conditions, supporting the reproducibility and robustness of this approach, we also uncovered numerous cell-type-specific and ADAR1-or ADAR2-specific interactors. Cell-type-specific hits may arise from differences in expression of those proteins between cell types, however a similar explanation would not explain ADAR-specific hits. We cannot rule out that some of the proteins specific to each condition may be an artifact of the common variability seen in mass-spectrometry-based screens; however, the reproducibility of the data suggests strong biological signals in the dataset. This dataset is highly complementary to and greatly extends a recently published study of RNA editing regulators identified through analysis of the ENCODE RNA binding protein knockdown RNA-seq dataset (Quinones-Valdez et al., 2019). We have uncovered 269 proteins that interact with ADARs, many of which overlap with the recently identified RNA binding proteins that regulate editing, validating our approach. Our biochemical screen enabled us to identify additional regulators, many of which are potentially ADAR and cell-type-specific.

When we first set out to identify *trans* regulators of editing, we were investigating two major potential mechanisms that could account for the observed tissue-specific differences in editing. *Trans* regulators could act directly on ADAR proteins to modify their activity at all sites equally but be differentially expressed in different tissues. Alternatively, *trans* regulators could affect a subset of sites by only interacting with or regulating ADAR proteins at those sites. We used publicly available RNA-seq and eCLIP-seq data from the ENCODE project to determine that we had uncovered both global and site-specific RNA editing regulators. Specifically, we found that the majority of our hits profiled by the ENCODE project showed binding at or near editing sites, and knocking down those proteins resulted in changes in the editing levels specifically at the sites they bound. This finding suggests that many of the editing sites regulated in *trans* are controlled by proteins interacting with ADAR proteins at editing sites. In some cases, the primary function may not be to regulate editing, but they nevertheless alter ADAR binding and editing. For example, BioID recovered many of the proteins found in paraspeckles, a complex ADAR1 had previously been found to be associated with (Anantharaman et al., 2016), highlighting the power of BioID to identify nearly all proteins found in a complex. We also found proteins that are ADAR interactors but not regulators of editing. These proteins may help ADAR proteins perform their other known functions in the cell such as miRNA regulation, or as yet unknown editing-independent functions of ADARs. They may also regulate the translation of ADARs, similar to PTBP1, which can activate the translation of ADAR-p110 through an IRES-like element (Yang et al., 2015).

The most intriguing finding of the BioID screens was the identification of all four DZF-domain containing proteins among the strongest hits. This finding suggests the importance of this class of proteins as ADAR interactors. These proteins largely interact with ADARs in an RNA-dependent fashion. ILF3 and STRBP appear to compete with ADAR to bind and edit dsRNAs because overexpression of these proteins led to decreased editing overall. In particular, ILF3 inhibited editing at a large number of sites, suggesting that it is a strong global suppressor of editing. This finding is consistent with a recent report that knockdown of ILF3 decreases editing at specific sites in K562 cells (Quinones-Valdez et al., 2019). Because ILF3 and ADAR2 have structurally similar dsRBDs and are able to compete for substrates in a biochemical assay (Wolkowicz and Cook, 2012) we tested whether this mechanism held in cells. We overexpressed a mutant version of ILF3 that lacks both dsRNA binding domains and found that the mutant did not interact with either ADAR and was unable to suppress editing. As we found that a number of RNA binding proteins bound RNA near the editing sites that they regulated, we further hypothesize that competing with ADAR for RNA binding is a prominent regulatory mechanism of RNA binding proteins on editing levels.

It will be interesting to more thoroughly explore the role of the other DZF domain containing proteins, ZFR and STRBP, in interacting with ADARs and regulating editing. The companion paper from our lab shows that Zinc finger RNA binding protein Zn72D, the fly homolog of ZFR, regulates over half of editing events in the fly brain, and that knockdown of ZFR leads to a decrease in a large number of editing events in mouse primary neurons. Zn72D also interacts with dADAR in an RNA-dependent manner, suggesting a similar mechanism to ILF3 regulation of ADARs that we detail here, but ZFR appears to be a positive regulator of editing, in contrast to ILF3. ZFR regulates editing of primarily ADAR2 editing sites in mouse primary neurons, suggesting that ZFR may be a particularly important regulator of ADAR2 editing in the brain (Sapiro et al., n.d.). Future work may explore whether STRBP has a similarly strong tissue-specific effect on editing, as it is highly expressed in the brain and testes. Together, our work suggests that DZF-domain-containing proteins as a class are critical for proper RNA editing, and future work is needed to further explore the mechanistic details and cell-type specificity of this role, including the potential role of the actual DZF-domain in supporting a protein interaction with ADAR.

The BioID experiments uncovered a large number of novel ADAR interactors and putative RNA editing regulators. These hits can be further characterized for their roles in both RNA editing and/or non-canonical roles for ADAR in the cell. Such knowledge may enable the manipulation of editing levels at specific sites without the manipulation of ADAR proteins themselves, which may have therapeutic benefit for cancer, autoimmune diseases, neurological diseases, or other diseases in which ADARs play a role.

## Methods

### Cell Culture, Transfections and Viral Transductions

HeLA S3 and 293T cells were cultured in DMEM supplemented with 10% FBS and Pen/Strep. M17 cells were cultured in F12 media supplemented with 10% FBS and Penn/Strep. Lentivirus was generated using standard methods: briefly, 293T cells were transfected with 3^rd^ generation packaging constructs and target plasmid for 8 hours then media was changed and viral supernatant was collected at 24 and 48 hour after and filtered at .45 uM. Cells were transduced with 1X viral supernatants supplemented with 5 ug/ml polybrene. Media was changed 12-24 hours later and selection began with the appropriate antibiotics between days 3-7.

BirA* constructs (BirA*-ADAR1, BirA*-ADAR2, BirA*-GFP) were created by subcloning the cDNA for each gene into the pCDH backbone containing a 3XFLAG tag, a nuclear localization sequence and the BirA* cDNA sequence upstream of the target gene, resulting in 3xFLAG-NLS-BirA*-ADAR1, 3xFLAG-NLS-BirA*-ADAR2, and 3xFLAG-NLS-BirA*-GFP constructs and, when translated, fusion proteins. These constructs were used to generate lentivirus, as described above.

FLAG-tagged constructs for ADARs (ADAR1 and ADAR2) and DZF-domain-containing proteins (ILF2, ILF3, STRBP, ZFR) were generated by subcloning the cDNA for each gene into the pCDH backbone containing an upstream (N-terminal) 3xFLAG. These constructs were used to generate lentivirus, as described above.

A non-tagged version of ADAR2 (for Figure 3) was generated by subcloning the ADAR2 cDNA into the pCDH backbone.

All transient transfection constructs were generated by subcloning the cDNA of interest into the pCDNA 3.1-3xFLAG (GFP, ADAR1, ADAR2, ILF2, ILF3, STRBP). All transient transfections were performed with Lipofectamine 2000 according to manufacturer’s protocol.

ILF3 ΔdsRBD was generated by synthesizing an ILF3 cDNA lacking the nucleotides coding for amino acids 402-572 (inclusive) and subcloning it into pCDNA 3.1-3xFLAG and pCDH.

### BioID Experimental Procedure

Each of the three (ADAR1, ADAR2, and GFP) BirA constructs were stably expressed in HeLa and M17 cells via lentiviral transduction. Ten 150mm plates of confluent cells expressing each construct were grown overnight in media with a final concentration of 50 uM D-Biotin (Life technologies, B-20656).

### Nuclear Lysate Preparation

Cells were harvested, pelleted and washed once with 1 X PBS. A 10X pellet volume of ice cold cytoplasmic extraction buffer (10 mM Hepes pH 7.5, 10 mM KCL, 1.5 mM MgCl2, 0.34 M Sucrose, 10% glycerol) with 1 mM DTT and protease inhibitors (cOmplete, Mini, EDTA-free Protease Inhibitor Cocktail Tablets, # 4693159001) was added and the pellet was gently resuspended. Cells were incubated on ice for 15 min. Triton X-100 was added to a final concentration of 0.1% and cells were vortexed for 10 seconds then incubated on ice for 5 min. Cells were spun down at 1300xg for 5 min at 4C, and washed once with cytoplasmic extraction buffer, spun down again and the supernatant was discarded. The nuclei were lysed with 7X the volume of the original pellet with high salt NP-40 lysis buffer (25 mM Hepes pH 7.5, 420 mM NaCl, 1.5 mM MgCl2, 10% glycerol, 0.5% NP-40) and incubated on ice for 20 min with occasional vortexing. Lysate was then spun at >20,000xg for 15 min at 4C. Protein concentration was then measured by BCA (Pierce, 23227).

### Preclear Lysate

1 mL of Agarose Control Resin (Pierce, 26150) slurry was added per sample to a 10 mL centrifuge column (Pierce, 89898) and placed inside a 50 mL conical tube and spun at 1000xg for 1 min to remove storage buffer. The resin was washed once with 4X resin bed volume of high salt NP-40 lysis buffer, and spun at 1000xg for 1 min. Nuclear lysate was added to the washed resin and incubated at 4C for >2hrs, with rotation. The resin was placed in a new 50 mL conical, and spin at 1000xg for 1 min to collect the precleared lysate. The protein concentration is measured by BCA (Pierce, 23227).

### Bind Biotinylated targets to Streptavidin beads

20 uL of High Capacity Streptavidin Agarose (Pierce, 20359) per 1 mg of lysate is added to a 10 ml centrifuge column placed inside a 50 mL conical tube. 6 volumes of high salt NP-40 lysis buffer is added to column and allowed to drain by gravity flow. Nuclear lysate is added to washed resin and incubated at 4 °C overnight, with rotation.

### Washing and Eluting

Eluate was collected by gravity flow and retained as the depleted fraction. The resin was then stringently washed twice with 10X volumes of the resin bed volume of high salt NP-40 lysis buffer with 0.3% SDS and then twice with high salt NP-40 lysis buffer with 1.0% SDS. The washed resin was resuspended in 900 ul of elution buffer (1XPBS, 5% SDS, 10 mM D-Biotin (Life technologies, B-20656), transferred to an eppendorf tube and boiled for 15 min. The resin is then placed in a 2 mL centrifuge column (Pierce, 89896) placed inside a 15 mL conical tube and spun at 1000xg for 1 min to collect the eluate. MeOH/Chloroform precipitation was used to concentrate the eluate. In brief, to 150 μL of eluate 600 μL of methanol was added followed by 150 μL of chloroform and vortexed. 450 μL of ultrapure water was added and vortexed then centrifuged at 14,000 xg for 5 min. Upper aqueous phase was removed and 450 μL of methanol, 1 uL GlycoBlue was added and vortexed. The protein was pelleted by centrifuging at 14,000xg for 5 min and then methanol removed completely. Pellets were resuspended in 25 ul of 1x SDS-PAGE sample buffer then boiled at 95C for 5 min.

### BioID Mass Spectrometry and Analysis

#### Reagents and Chemicals

Deionized water was used for all preparations. Buffer A consists of 5% acetonitrile 0.1% formic acid, buffer B consists of 80% acetonitrile 0.1% formic acid, and buffer C consists of 500 mM ammonium acetate 0.1% formic acid and 5% acetonitrile.

#### Sample Preparation

Proteins were precipitated in 23% TCA (Sigma-Aldrich, St. Louis, MO, Product number T-0699) at 4 °C O/N. After 30 min centrifugation at 18000xg, protein pellets were washed 2 times with 500 ul ice-cold acetone. Air-dried pellets were dissolved in 8 M urea/ 100 mM Tris pH 8.5. Proteins were reduced with 5 mM Tris(2-carboxyethyl)phosphine hydrochloride (Sigma-Aldrich, St. Louis, MO, product C4706) and alkylated with 55 mM iodoacetamide (Sigma-Aldrich, St. Louis, MO, product I11490). Proteins were digested for 18 hr at 37 °C in 2 M urea 100 mM Tris pH 8.5, 1 mM CaCl_2_ with 2 ug trypsin (Promega, Madison, WI, product V5111). Digest was stopped with formic acid, 5% final concentration. Debris was removed by centrifugation, 30 min 18000xg.

#### MudPIT Microcolumn

A MudPIT microcolumn(4) was prepared by first creating a Kasil frit at one end of an undeactivated 250 □m ID/360 □m OD capillary (Agilent Technologies, Inc., Santa Clara, CA). The Kasil frit was prepared by briefly dipping a 20 - 30 cm capillary in well-mixed 300 □L Kasil 1624 (PQ Corporation, Malvern, PA) and 100 □L formamide, curing at 100°C for 4 hrs, and cutting the frit to ~2 mm in length. Strong cation exchange particles (SCX Luna, 5 □m dia., 125 Å pores, Phenomenex, Torrance, CA) was packed in-house from particle slurries in methanol 2.5 cm. An additional 2.5 cm reversed phase particles (C18 Aqua, 3 µm dia., 125 Å pores, Phenomenex) were then similarly packed into the capillary using the same method as SCX loading, to create a biphasic column. An analytical RPLC column was generated by pulling a 100 □m ID/360 □m OD capillary (Polymicro Technologies, Inc, Phoenix, AZ) to 5 □m ID tip. Reversed phase particles (Aqua C18, 3 □m dia., 125 Å pores, Phenomenex, Torrance, CA) were packed directly into the pulled column at 800 psi until 12 cm long. The MudPIT microcolumn was connected to an analytical column using a zero-dead volume union (Upchurch Scientific (IDEX Health & Science), P-720-01, Oak Harbor, WA).

LC-MS/MS analysis was performed using an Eksigent nano lc pump and a Thermo LTQ-Orbitrap using an in-house built electrospray stage. MudPIT experiments were performed where each step corresponds to 0, 20, 50 and 100% buffer C being run for 3 min at the beginning of each gradient of buffer B. Electrospray was performed directly from the analytical column by applying the ESI voltage at a tee (150 □m ID, Upchurch Scientific). Electrospray directly from the LC column was done at 2.5 kV with an inlet capillary temperature of 275 °C. Data-dependent acquisition of MS/MS spectra with the LTQ -Orbitrap were performed with the following settings: MS/MS on the 10 most intense ions per precursor scan, 1 microscan, reject charge unassigned charge state; dynamic exclusion repeat count, 1, repeat duration, 30 second; exclusion list size 200; and exclusion duration, 15 second.

#### Data Analysis

Protein and peptide identification and protein quantitation were done with Integrated Proteomics Pipeline - IP2 (Integrated Proteomics Applications, Inc., San Diego, CA. http://www.integratedproteomics.com/). Tandem mass spectra were extracted from raw files using RawExtract 1.9.9(1) and were searched against Uniprot human database with reversed sequences using ProLuCID(2, 5). The search space included half- and fully-tryptic peptide candidates. Carbamidomethylation (+57.02146) of cysteine was considered as a static modification. Biotinylation of lysine (226.077598) was considered as a variable modification, Peptide candidates were filtered using DTASelect, with these parameters -p 2 -y 2 --trypstat –pfp 0.01 --extra --pI -DM 10 --DB --dm -in -m 1 -t 1 --brief --quiet (1, 3).

### Analysis of BioID hits

BirA*-ADAR1 and BirA*-ADAR2 experiments were analyzed separately. In both cases, peptide counts were summed to generate protein-level counts. For each biological replicate, the fold change of the ADAR condition vs the no-BirA* and GFP conditions were separately calculated (1 was added to all counts in order to avoid infinite fold changes). Proteins were then filtered according to the following criteria: for each replicate, in order to be retained, a protein was required to have a fold change >2 versus both the no-BirA* and GFP conditions. A second level of filtering then required each protein to be retained in at least 2 out of the 3 biological replicates. These retained proteins were considered hits for that condition. For heatmaps, log_2_ fold change for the ADAR condition was calculated by determining the median log_2_ fold change versus the GFP condition across all three replicates.

### Immunoprecipitation

FLAG-tagged proteins were immunoprecipitated from 2 mg of HEK293T or M17 cells using anti-FLAG M2 affinity gel (Sigma A2220). Lysates were prepared in NP40 buffer (see Western blotting). 40 ul of the anti-FLAG M2 affinity resin was washed three times with 1 ml of lysis buffer then added to the 2 mg of protein extract diluted to 500 ul in NP-40 lysis buffer and incubated for 2 hours at 4°C with rotation. The resin was then washed 4 times for 5 minutes each with 1 mL NP-40 lysis buffer at 4°C with rotation. For further analysis the resin was resuspended with 30 ul 2x SDS-PAGE sample buffer, then incubated at 95 degrees for 5 min to elute bound proteins.

### Western Blotting

Cells were lysed in NP-40 buffer (25 mM HEPES-KOH, 150 mM KCl, 1.5 mM MgCl2, 0.5% NP40, 10% Glycerol [pH 7.5]) supplemented with protease inhibitors (cOmplete, Mini, EDTA-free Protease Inhibitor Cocktail Tablets, # 4693159001) Lysates were clarified by spinning for 10 min at 13,000 rpm at 4°C, and the protein content was measured by BCA (Pierce, 23225). 10 ug of protein was separated by SDS-PAGE, transferred onto nitrocellulose membrane, and blotted according to standard protocols. Chemiluminescence was imaged using a BioRad ChemiDoc imaging system.

### Antibodies

ILF2: Bethyl laboratories NF45 Antibody, cat: A303-147A (1:1000)

ILF3: Bethyl laboratories NF90 Antibody, cat: A303-651A (1:1000)

ZFR: Abcam Anti-ZFR antibody ab90865 (1:500)

STRBP: Abcam Anti-STRBP antibody cat: ab111567 (1:500)

ADAR1: Santa Cruz ADAR1 Antibody (15.8.6) cat: sc-73408 (1:500)

ADAR2: Genetex ADAR2 antibody cat: GTX114237(1:500)

FLAG: Sigma Monoclonal ANTI-FLAG® M2-Peroxidase (HRP) (1:5000)

### mmPCR-seq of samples

The mmPCR–seq was performed as described in (Zhang et al., 2014). Briefly, total RNA is extracted from cells using a Qiagen Micro or Mini RNeasy kit (Qiagen cat:74004 or 74104) and reverse transcribed using iScript Advanced reverse transcriptase (Bio-Rad). The cDNAs were purified using Ampure XP Beads (Beckman Coulter), with an elution volume of 10 ul. 250 ng of cDNA was used in a preamplification reaction and amplified cDNA was purified using Ampure XP beads and eluted in 10 ul. 50 ng of pre-amplified cDNA was loaded into each well of an Access Array microfluidic chip (Fluidigm). The PCR reactions were performed on the Access Array System (Fluidigm) using KAPA2G 5X Fast Multiplex PCR Mix (Kapa Biosystems). Barcodes were added in a second round of PCR using Phusion DNA polymerase (NEB cat: M0531S). Samples were sequenced with 76 base-pair paired-end reads using an Illumina NextSeq (Illumina, San Diego, CA).

### Analysis of mmPCR-seq and RNA-seq

mmPCR-seq and RNA-seq reads were mapped to the human genome (hg19) using STAR (Dobin et al., 2013) version 2.4.2a using the parameters (--outFilterMultimapNmax 20 -- outFilterMismatchNmax 999 --outFilterMismatchNoverReadLmax 0.1 --alignIntronMin 20 -- alignIntronMax 1000000 --alignMatesGapMax 1000000 --alignSJoverhangMin 8 -- alignSJDBoverhangMin 1 --sjdbScore 1 --twopassMode Basic). For editing analysis we used the Samtools (version 0.1.16) (Li et al., 2009) mpileup command to count A and G counts at known editing sites (Tan et al., 2017), RADAR version 2 (Ramaswami and Li, 2014). For RNA-seq only bases with quality scores > 20 were used. For both RNA-seq and mmPCR-seq, combined A and G counts from two replicates of each condition were compared to down sampled GFP overexpression reads using Fisher’s exact test with a Benjimini-Hochberg multiple hypothesis testing correction in R (Benjamini and Hochberg, 1995).

For both RNA-seq and mmPCR-seq analysis, any variability between replicates could arise for a number of reasons, e.g. low expression or inefficient primer-based amplification, so for all subsequent analyses we only assessed sites with high coverage (50X in mmPCR-seq and 20X in RNA-seq) that were <10% different between replicates (**Figure S6A, B**).

For gene expression analysis, FPKMs were calculated using RSEM 1.2.21 (Li and Dewey, 2011). To compare global expression differences of cell-type-specific BioID hits in M17 and HeLa cells, we used Wilcoxon matched-pairs signed ranked test performed by Graphpad PRISM 7.

### RNA-seq library preparation

Total RNA was extracted from cells using Qiagen Micro or Mini RNeasy kit (Qiagen cat:74004 or 74104). rRNA was depleted from total RNA following RNase H-based protocols adopted from (Adiconis et al., 2013; Morlan et al., 2012). We mixed approximately 250 ng of RNA with 61.54 pmoles of pooled DNA oligos designed antisense to rRNA (gift from J. Salzman lab at Stanford) in a 5ul reaction with 1 ul of 5X RNase H(-)Mg buffer (500 mM Tris-HCl pH 7.4, 1 M NaCl) and 0.25 ul 1mM EDTA. We annealed rRNA antisense oligos to total RNA samples for 2 minutes at 95°C, slowly reduced the temperature to 45°C and then added 5 ul of RNase H mix (1.7 ul water, 1 ul 5X RNase H(-)Mg buffer, 0.2 ul 1 M MgCl_2_, 0.1ul RiboLock RNase Inhibitor (40 U/µL) (ThermoFisher EO038), 2 ul of Hybridase Thermostable RNase H (Epicenter, Madison, WI: Lucigen H39500) to make 10 ul total and incubated for 30 minutes at 45°C. rRNA-depleted RNA was then purified using 2.2X reaction volume of Agencourt RNAClean XP beads (Beckman Coulter: A63987), treated with TURBO DNase (Invitrogen: AM1907), and then purified with RNAClean XP beads again. rRNA-depleted RNA was used as input to the KAPA HyperPrep RNA-seq Kit (Kapa Biosystems: KK8540). All libraries were sequenced with 76 base-pair paired-end reads using an Illumina NextSeq (Illumina, San Diego, CA).

### Sanger Sequencing of RNA levels

To determine editing levels at PAICS and GRIA2 editing sites, RNA was extracted from M17 cells expressing BirA*-GFP, BirA*-ADAR1, and BirA*-ADAR2 using Zymo Quick RNA kit. RNA was treated with TURBO DNase and then cDNA was synthesized using Bio-Rad iScript Advanced cDNA synthesis kit. PCR was performed using the following to amplify around the editing site in each gene, PAICS FWD: TCAATCCACCCTTTTCCAAG, REV: TGATAAAAACGTGGGCCTTC and GRIA2 FWD: CAGCAGATTTAGCCCCTACG REV: AGATGAGATGTGTGCCAACG) with NEB Phusion for 40 cycles, and amplicons were gel purified with Qiagen QIAquick Gel Extraction kit (Cat 28115) and sent for Sanger sequencing. Two replicates were performed for each group of cells.

### ENCODE data analysis

We downloaded the mapped BAM files (HG38 version) of RNA-seq data generated following RBP knockdown or control shRNA transfection from the ENCODE data portal (encodeproject.org). For the RNA-seq data, we used the same pipeline as described in the Analysis of mmPCR-seq and RNA-seq section to quantify editing levels. Similarly, as reported (Quinones-Valdez et al., 2019), we also found batch effects in the editing level measurements from the RNA-seq data. However, in this paper we only performed editing level comparison between RBP knockdown and the matched control shRNA transfection within the same batch, so it was not necessary to remove the batch effects that only affected editing level measurements of samples from different batches. When the accumulative editing level differences was calculated and compared between different RBPs in **Fig. 2E**, the influence of batch effects was taken into consideration by normalizing the CDED to [0,1].

To determine where the RBPs bind on the RNA, we analyzed the eCLIP-seq data from ENCODE. We downloaded the BED files (HG38 version) of the called CLIP peaks for each eCLIP-seq data and used shifted z-score method to test how the strength of association between the RBP binding and the editing sites would change if the peak was shifted from its original position (Gel et al., 2016). The z-scores were calculated as the distance between the expected binding value and the observed one, measured in standard deviations. And we shifted the peaks 50bp stepwise within the 1.2kb up- and down-stream windows of the editing site to obtain the corresponding z-score values.

## Accession Numbers

The high-throughput sequencing data utilized in this work, including the RNA-seq and mmPCR-seq libraries, have been deposited in the Gene Expression Omnibus (GEO) database, accession number GSE130771.

## ACKNOWLEDGEMENTS

We thank Adam Freund, and all members of the Li Lab especially Tao Sun for useful discussions and comments on this paper. We would also like to thank Julia Salzman for her generous donation of the oligos used for rRNA depletion in the RNA-seq library preparation. Funding sources: American Heart Association Postdoctoral grant 16POST27700036 (to E.C.F), National Center for Research Resources 5P41RR011823 (to JJM and JRY), NIH R01-GM102484 (to J.B.L.), R01-GM124215 (to J.B.L.), R01-MH115080 (to J.B.L.)

---

**Figure S1.**
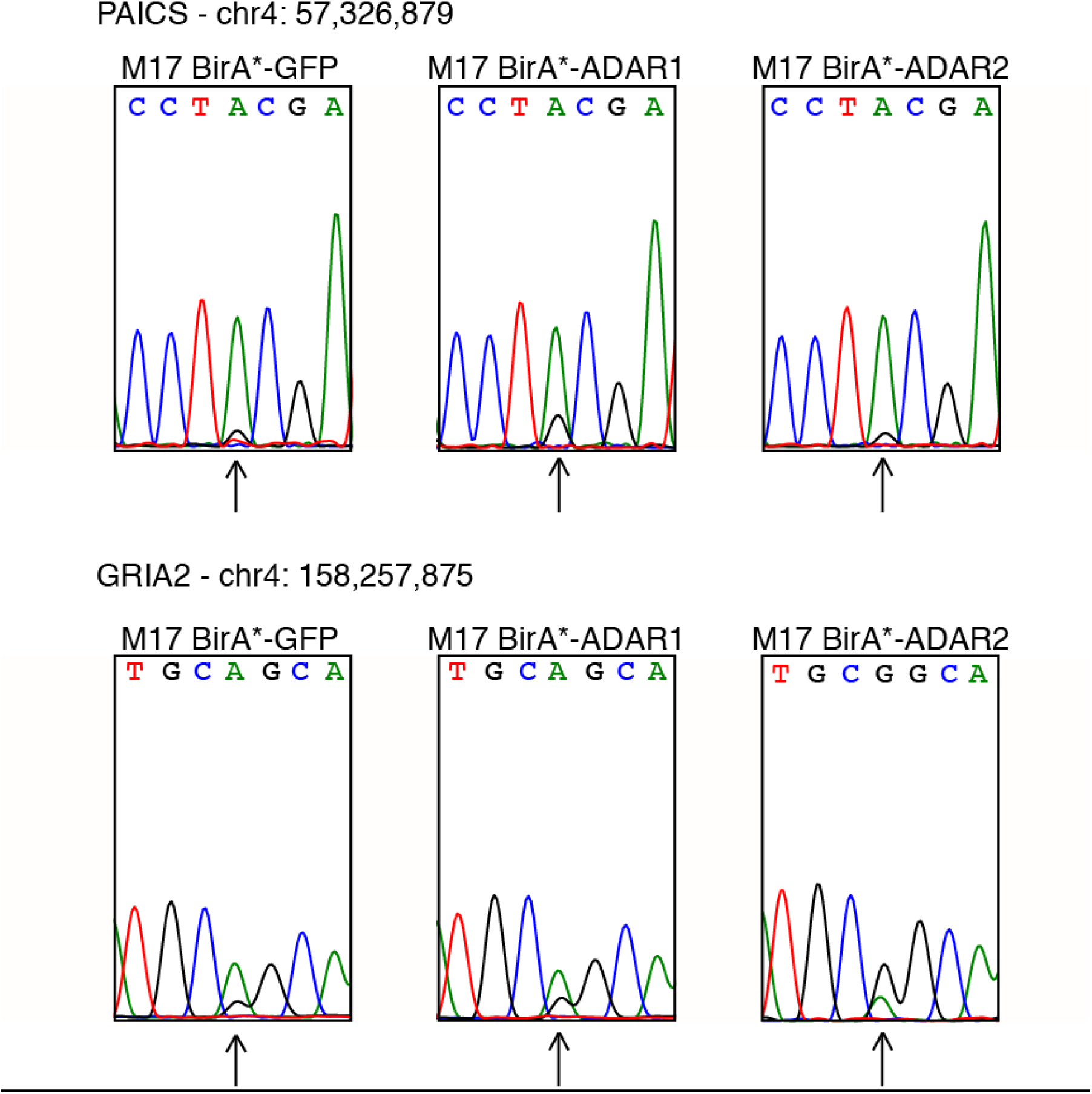
BirA*-ADAR1 and BirA*-ADAR2 retain editing activity. (A) Sanger sequencing traces at an ADAR1-regulated editing site in *PAICS* (chr4: 57,326,879) in M17 cells expressing BirA*-GFP, BirA*-ADAR1, and BirA*-ADAR2 assayed in BioID experiments. The site is most highly edited in cells expressing BirA*-ADAR1. (B) Sanger sequencing traces at an ADAR2-regulated editing site in *GRIA2* (chr4: 158,257,875) in M17 cells expressing BirA*-GFP, BirA*-ADAR1, and BirA*-ADAR2 assayed in BioID experiments. The site is most highly edited in cells expressing BirA*-ADAR2. Arrows denote each editing site in the sequence.

**Figure S2.**
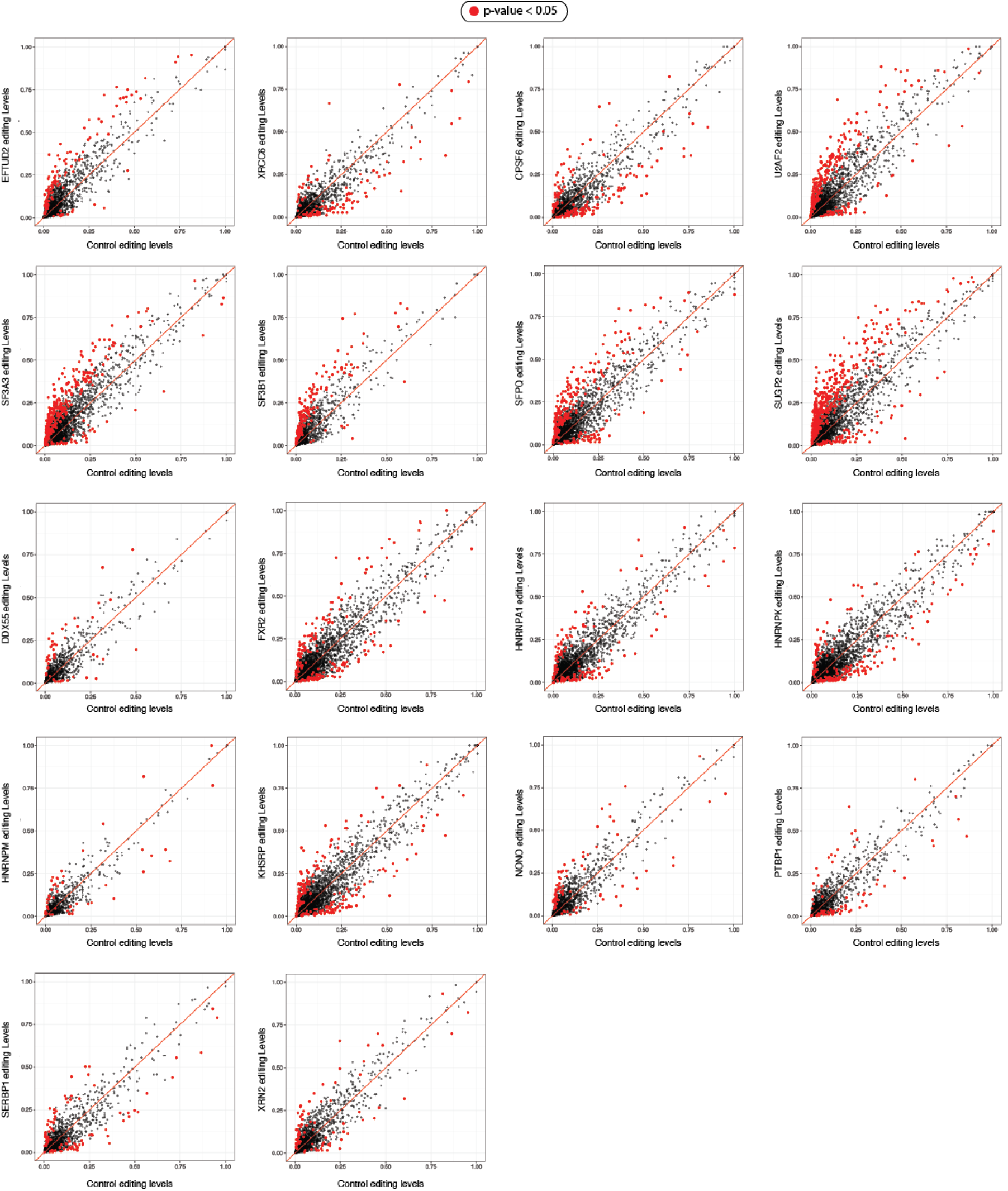
Scatter plots of pairwise comparison of editing levels between knockdown and control RNA-seq of RBPs that were also found in the BioID assay. Red dots, Fisher’s exact test p-value < 0.05.

**Figure S3.**
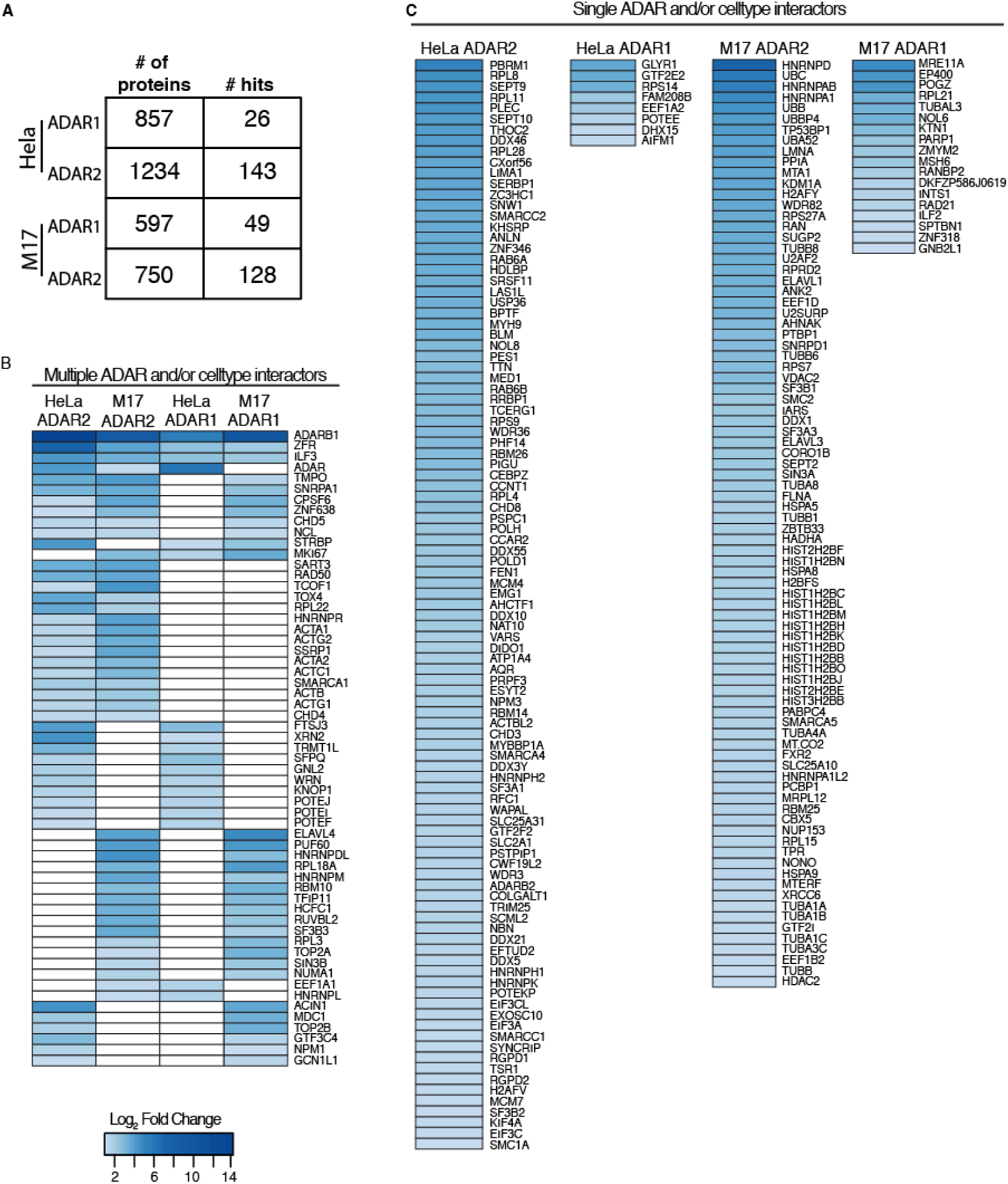
BioID of ADAR1 and ADAR2 in HeLa and M17 cells reveals interactors specific to each cell type and each ADAR. (A) The total number of proteins identified by mass spec from each IP and the number of hits remaining after the filtering pipeline illustrated in Figure 1C. (B) A heatmap displaying the hits identified in multiple IPs. All proteins detected in at least two conditions are displayed, arranged by unsupervised hierarchical clustering. The strength of blue indicates the log fold change over the GFP control. White indicates that the protein was not detected in the IP. (C) A heatmap displaying all proteins identified in only one condition. The strength of blue indicates the log fold change over the GFP control.

**Figure S4.**
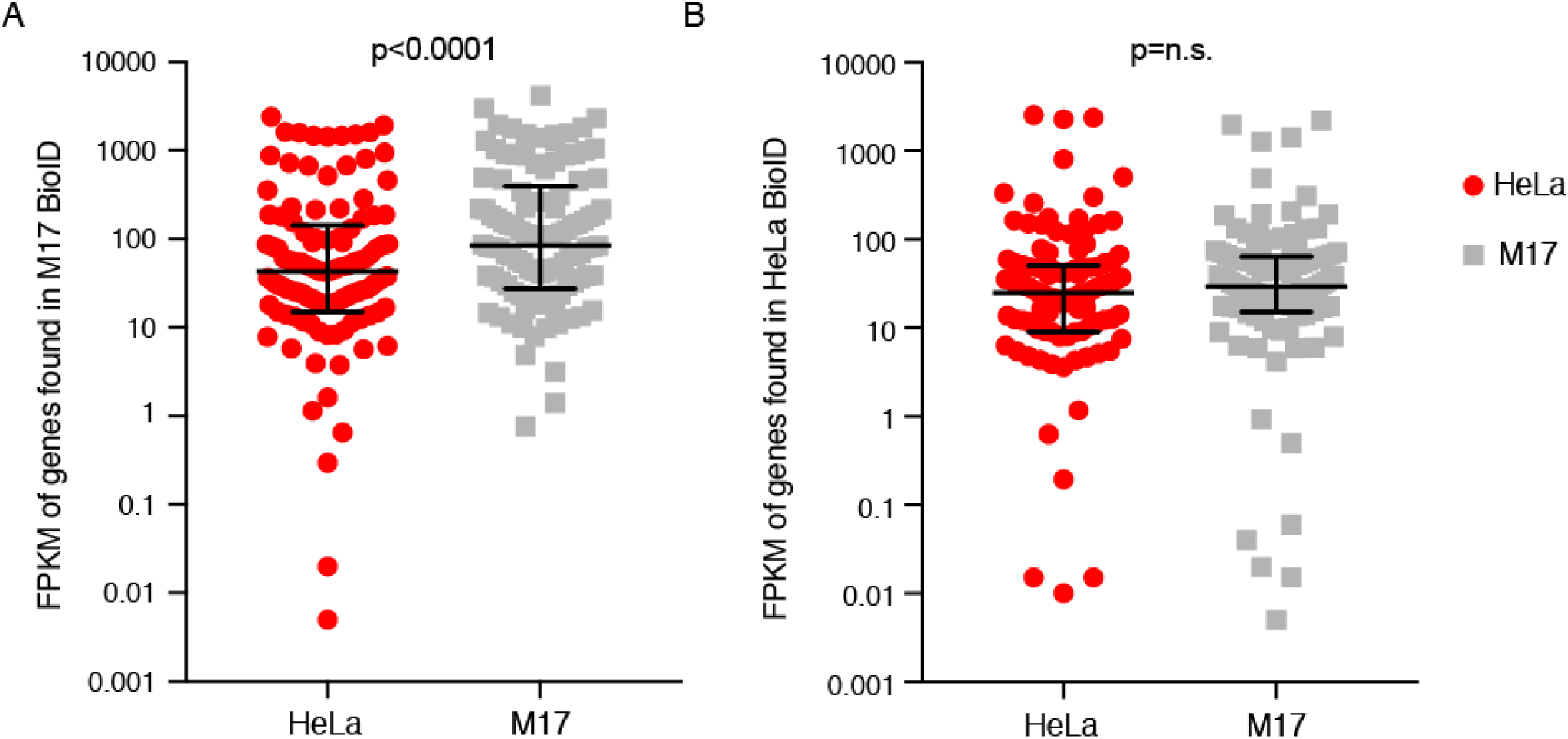
Genes found in the M17 BioID are more highly expressed in M17 cells. **A.** Gene expression of all genes that encode proteins identified in the M17 ADAR1 and ADAR2 BioID screens. Each dot represents the FPKM in HeLa (red) or M17 (grey) cells, as assayed by RNA-seq. Overall, the set of genes is more highly expressed in M17 cells versus HeLa. **B.** Gene expression of all genes that encode proteins that were identified in the HeLa ADAR1 and ADAR2 BioID screens. Each dot represents the FPKM in HeLa (red) or M17 (grey) cells, as assayed by RNA-seq. There is not a significant difference between HeLa and M17. The median (middle black bar) with interquartile range (HeLa, black bars and M17, black bars) are shown for each plot. P-values were determined by Wilcoxon matched pairs signed rank test.

**Figure S5.**
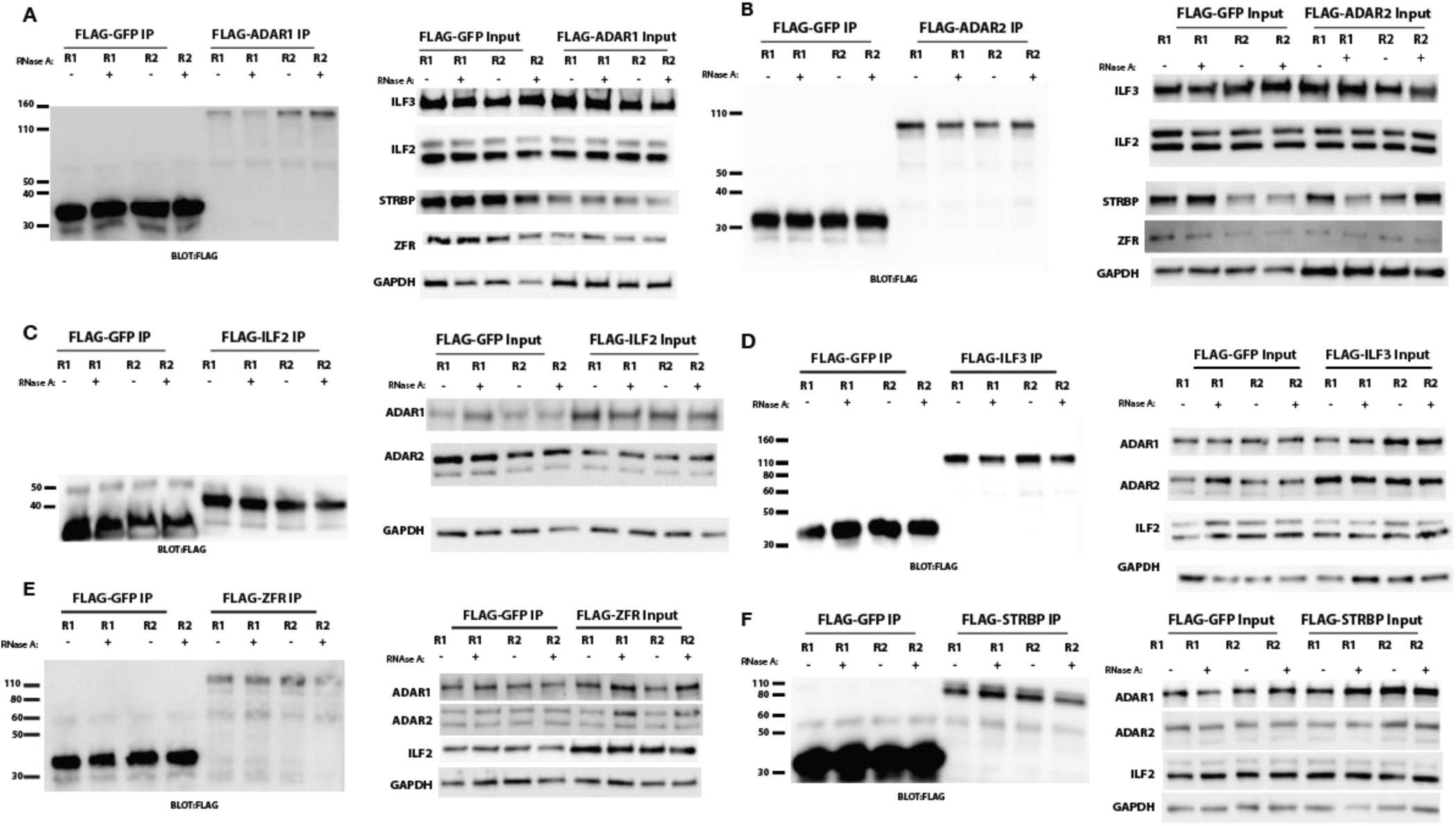
Input of the IPs from Figure 3 show strong expression of each FLAG-tagged construct and blotted protein. **A.** Inputs of IPs from Figure 3B. **B.** Inputs of IPs from Figure 3C. **C.** Inputs of IPs from Figure 3D. **D.** Inputs of IPs from Figure 3E. **E.** Inputs of IPs from Figure 3F. **F.** Inputs of IPs form Figure 3G.

**Figure S6.**
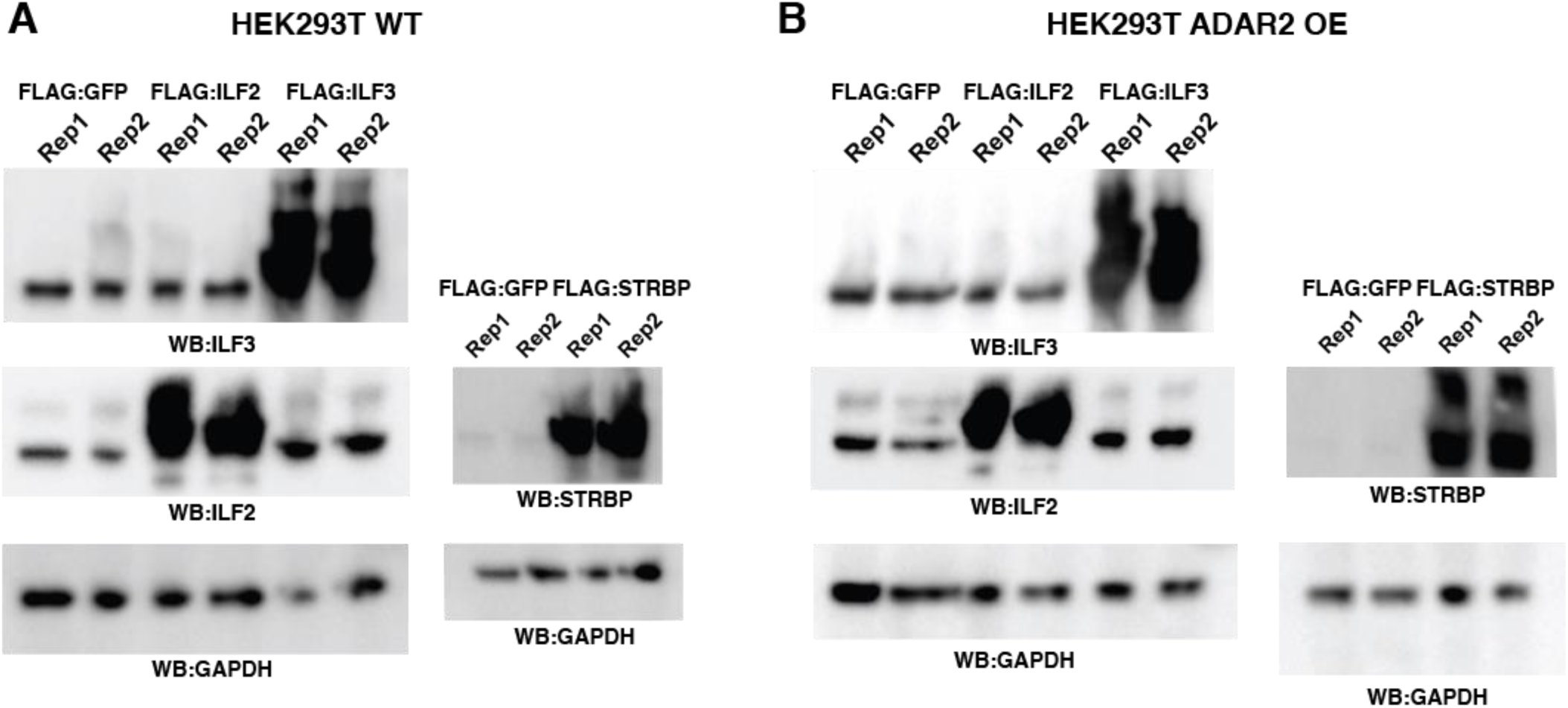
The overexpression of each DZF-domain-containing protein in the cells lines assayed by RNA-seq and mmPCR-seq in Figure 3 were determined by western blot. **A**. Protein from HEK293T cells transiently overexpressing FLAG fused to ILF2, ILF3 (left) and STRBP (right) was blotted for ILF3, ILF2, STRBP and GAPDH as a control. **B.** Protein from HEK293T A2 OE cells transiently overexpressing FLAG fused to ILF2, ILF3 (left) and STRBP (right) was blotted for ILF3, ILF2, STRBP and GAPDH as a control.

**Figure S7.**
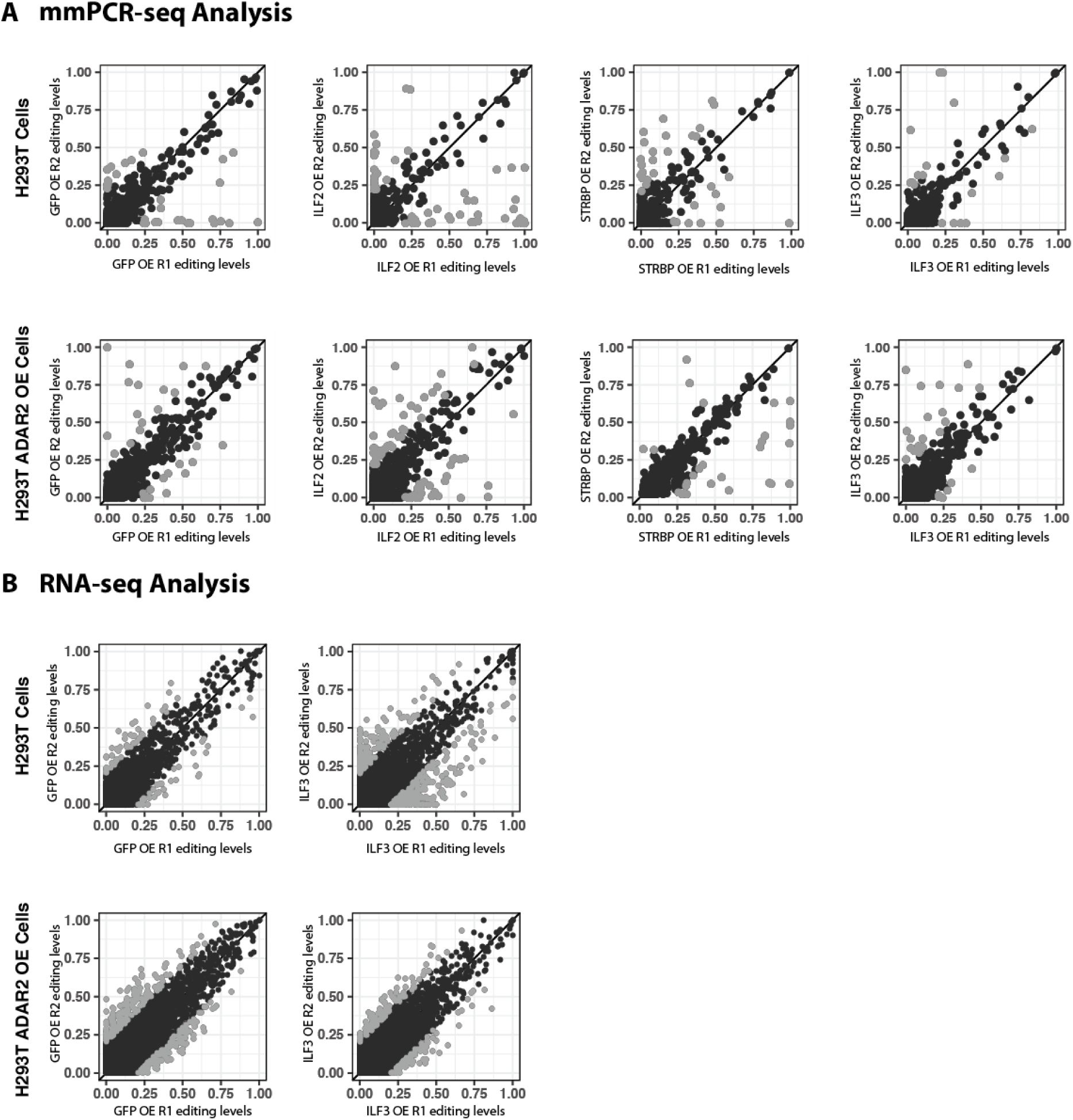
mmPCR-seq to measure editing levels demonstrates low variability in biological replicates. Sites with high variability were removed from further analysis. **A**. Scatterplots of biological replicates assayed by mmPCR-seq from HEK293T (top row) and HEK293T ADAR2 OE (bottom row) overexpressing GFP, ILF2, STRBP or ILF3 (from Figure 3). Gray dots represent sites with more than 20% variability between replicates, and were not included in further analysis. **B.** Scatterplots of biological replicates assayed by RNA-seq from HEK293T (top row) and HEK293T ADAR2 OE (bottom row) overexpressing GFP (left) or ILF3 (right). Gray dots represent sites with more than 20% variability between replicates, and were not included in further analysis.

**Figure S8.**
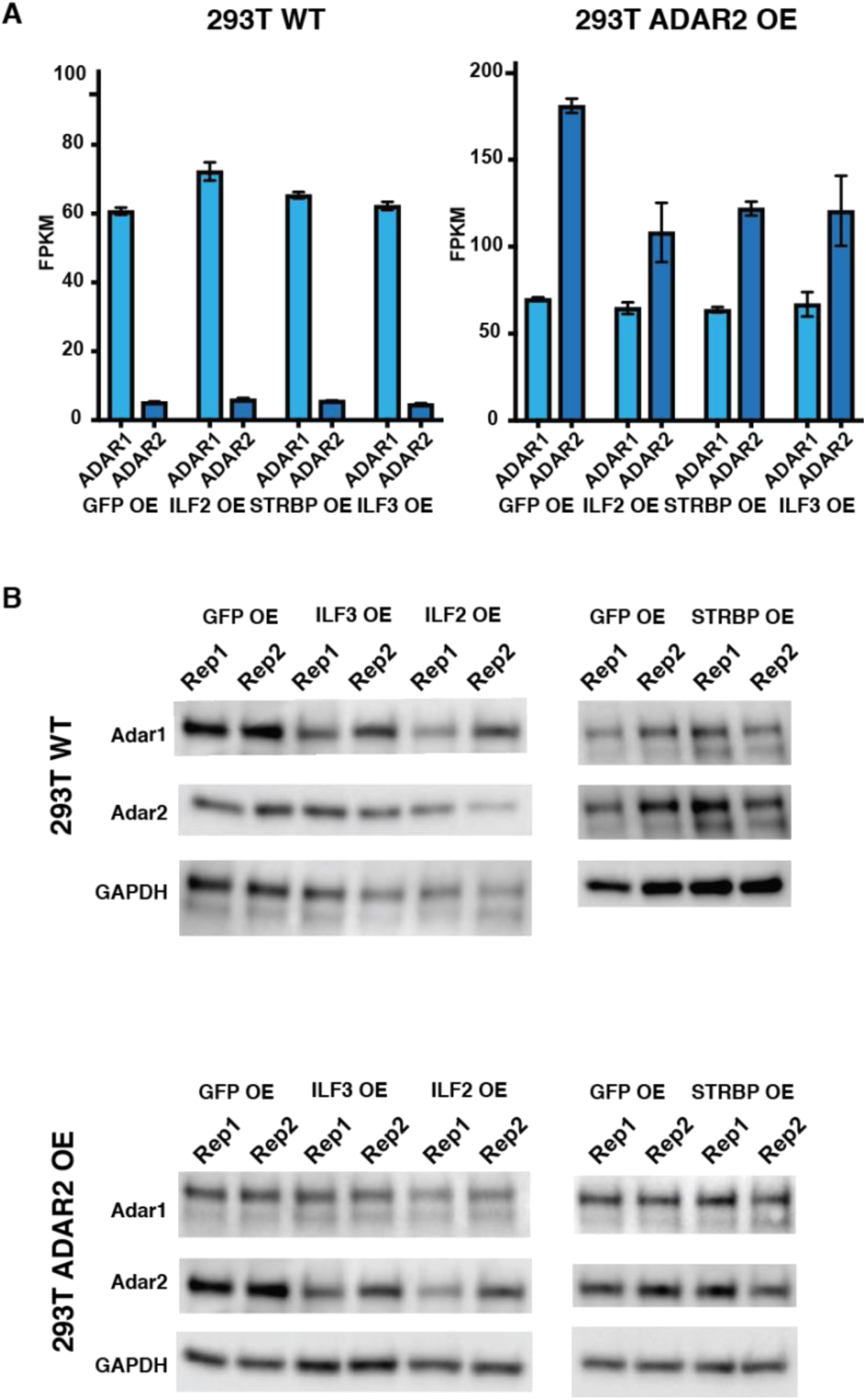
Levels of ADAR1 and ADAR2 are not greatly changed in cells lines overexpressing DZF-domain-containing proteins. **A.** Levels of ADAR1 (light blue) and ADAR2 (dark blue) in HEK293T WT (left) and HEK293T A2 OE (right) cells overexpressing GFP, ILF2, STRBP or ILF3. n = 2 biological replicates. The transcript levels of ADAR1 are not significantly changed in HEK293T or HEK293T A2 OE cells. The transcript levels of ADAR2 are reduced in HEK293T A2 OE cells overexpressing DZF-domain-containing proteins compared to GFP. **B.** Protein from HEK293T (top) or HEK293T A2 OE (bottom) cells overexpressing GFP, ILF2, ILF3 or STRBP were blotted for ADAR1, ADAR2 and GAPDH as a control. The protein levels of ADAR1 and ADAR2 are not greatly changed in wildtype or HEK293T A2 OE cells.

**Figure S9.**
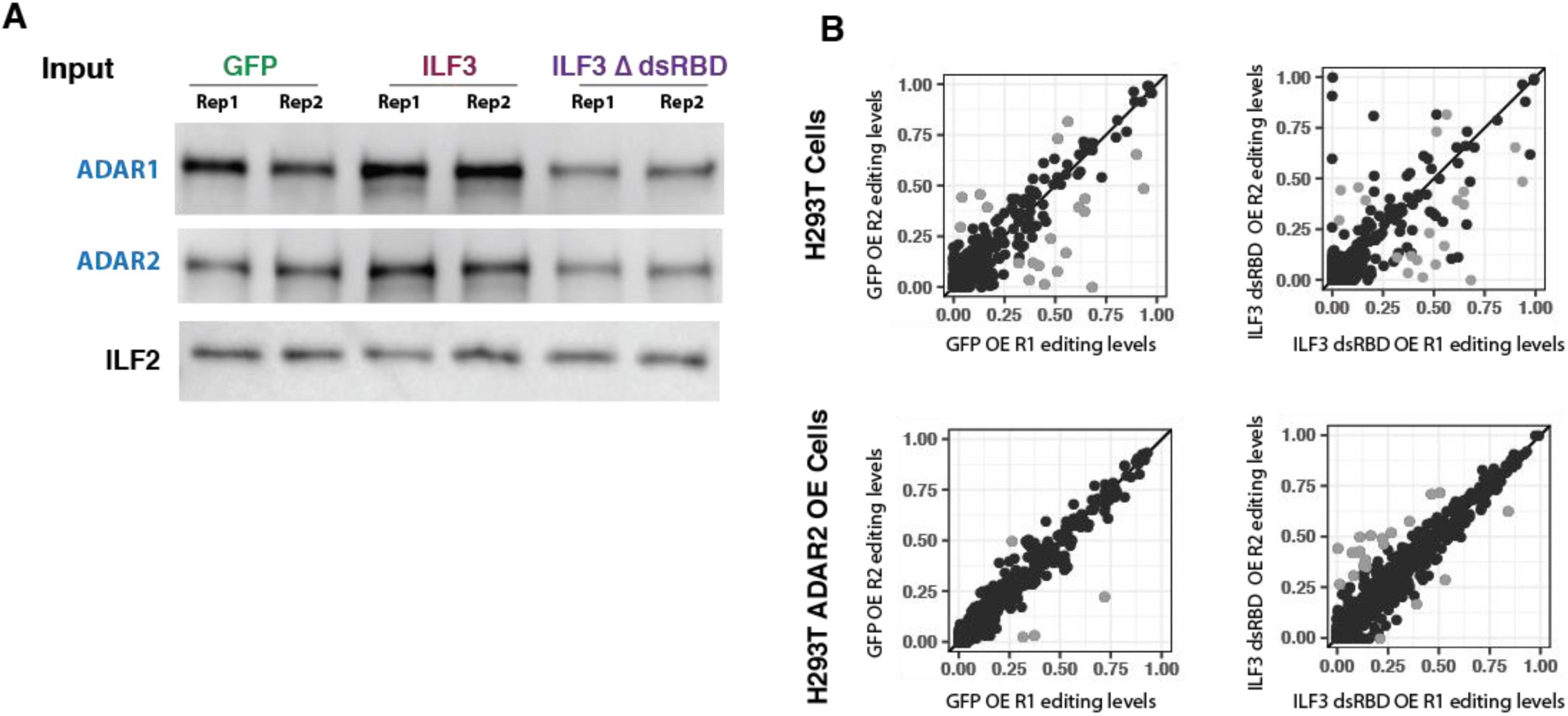
Biological replicates of cells overexpressing ILF3 and ILF3 dsRBD show consistent editing levels. **A.** Scatterplots of biological replicates assayed by mmPCR-seq from HEK293T (top row) and HEK293T A2 OE (bottom row) overexpressing GFP or ILF3 dsRBD (from Figure 5C). **B.** Inputs of IPs from Figure 5D.

## LIST OF SUPPLEMENTARY TABLES

**Supplementary Table 1. mmPCR-seq of HEK293T and HEK293T Adar2 overexpression with overexpression of ILF2, ILF3, ILF3dsRBD, and STRBP editing levels and p-values for comparison with GFP overexpression controls.**

**Supplementary Table 2. RNA-seq of HEK293T and HEK293T Adar2 overexpression with ILF3 overexpression editing levels and p-values for comparison with GFP overexpression controls.**

